# Targeting fatty acid synthase suppresses tumor development in *NF2/CDKN2A*-deficient malignant pleural mesothelioma

**DOI:** 10.1101/2024.07.14.603191

**Authors:** Sivasundaram Karnan, Akinobu Ota, Muhammad Nazmul Hasan, Hideki Murakami, Md. Lutfur Rahman, Md Wahiduzzaman, Md Towhid Ahmed Shihan, Nushrat Jahan, Lam Quang Vu, Ichiro Hanamura, Akihito Inoko, Miho Riku, Hideaki Ito, Yoshifumi Kaneko, Yinzhi Lin, Toshinori Hyodo, Hiroyuki Konishi, Shinobu Tsuzuki, Yoshitaka Hosokawa

**Affiliations:** Department of Biochemistry, Aichi Medical University School of Medicine, Nagakute, Aichi, Japan; Department of Food and Nutritional Environment, College of Human Life and Environment, Kinjo Gakuin University, Nagoya, Aichi, Japan; EuGEF Research Foundation, Chattogram, Bangladesh; Department of Pathology, Aichi Medical University School of Medicine, Nagakute, Aichi, Japan; Department of Biochemistry, Emory University School of Medicine, Atlanta, GA, USA; Department of Foundations of Medicine, NYU Grossman Long Island School of Medicine, 101 Mineola Blvd, Mineola, NY-11501, United States; Division of Hematology, Department of Internal Medicine, Aichi Medical University School of Medicine, Nagakute, Aichi, Japan; Department of Microbiology and Immunology, Aichi Medical University School of Medicine, Nagakute, Japan

**Keywords:** CRISPR/Cas9, FASN, p-Drp1, cerulenin, MPM

## Abstract

Malignant pleural mesothelioma (MPM) is an uncommon yet deadly cancer linked to asbestos exposure. The lack of effective early diagnosis and treatment leads to reduced life expectancy among patients with MPM. This study is aimed to identify a novel molecular target inhibitor to develop more effective therapeutics for MPM. Our drug screening assay showed that the fatty acid synthase (FASN) inhibitor cerulenin demonstrates strong and selective anti-proliferative properties against *NF2/CDKN2A(p16)*-deficient MPM cells, surpassing the effects of cisplatin or pemetrexed. FASN protein is frequently detected in *NF2/p16*-deficient MPM tumor-derived tissues (15/15, 100%), but rarely in *NF2/p16*-intact MPM tumors (8/25, 32%). Notably, cerulenin administration successfully reduced the growth of *NF2/p16*-deficient MPM tumors in xenografted mice. Cerulenin inhibits mitochondrial fission by targeting dynamin-related protein 1 (DRP1) in *NF2/p16*-deficient cells. Moreover, the disruption of the FASN gene leads to increased ubiquitination of DRP1. These findings suggest that FASN might play a role in the tumorigenesis of MPM cells through the regulation of mitochondrial dynamics. This research offers a novel perspective on the potential development of precision medicine for MPM.

## Introduction

Malignant pleural mesothelioma (MPM) is a highly aggressive neoplasm originating from pleural mesothelial cells, commonly linked to asbestos exposure after a latency period of 30–40 years (1–3). Diagnosis of MPM typically occurs in advanced stages, contributing to a grim prognosis for patients (4). MPM generally responds poorly to radiation and conventional chemotherapy (5). Recent advancements in the management of the disease, including diagnosis, staging biomarkers, and treatment strategies have provided greater understanding of MPM; however, the mortality rate remains high, partially due to late diagnosis and treatment resistance (6, 7).

Recent studies have revealed that MPMs are associated with frequent genetic alterations in the neurofibromatosis 2 (*NF2*), cyclin-dependent kinase inhibitor 2A (*CDKN2A*, *p16*), and BRCA1-associated protein 1 (*BAP1)* tumor suppressor genes (8–11). Previously, we have observed high expression levels of *FGFR2*, *CD24,* and *CAMK2D* in *NF2-*knockout (KO), *NF2/p16*-double KO (DKO), and *BAP1*-KO mesothelial cell lines, respectively (12–14), suggesting representation of diagnostic and therapeutic targets for MPM. However, a library containing 364 anticancer drugs was screened using *NF2/p16*-DKO human mesothelial cells to precisely understand the molecular determinants of MPM, and cerulenin—a fatty acid synthase (FASN) inhibitor was identified.

FASN—a key catalytic enzyme regulating long-chain fatty acid synthesis—is highly expressed in several human cancers, including MPM (15–17). FASN activity is necessary for mitochondrial priming (18) and senescence in cancer cells (19). FASN amplifies mitochondrial ATP synthesis (20) and promotes mitochondrial fission, key processes in the reprogramming of fatty acid metabolism in colon cancer cells (21). Mitochondria are highly pleomorphic and considered primary mediators of intracellular energy production (22, 23). Alterations in mitochondrial dynamics are associated with disease progression and drug resistance in various cancers (24–27); however, the precise role of FASN in mitochondrial biology in MPM cells remains unknown.

Inhibiting FASN activity prevents cancer cell proliferation, migration, invasion, cell cycle, signaling pathway, and energy metabolism in breast and colon cancers, diffuse large B-cell lymphomas, melanomas, retinoblastomas, prostate cancers, and mesothelioma cells (17, 28–35). Cerulenin, which is available as a naturally occurring or pharmacologically synthesized antibiotic, is an effective apoptosis inducer, with mitochondria being the key player in cerulenin-mediated apoptosis (36). This occurs by disrupting the mitochondrial membrane and dysregulating the mitochondrial membrane proteins (37). The preferential activity of FASN and its inhibitor cerulenin are attractive targets for anticancer therapy; however, the underlying mechanism of cerulenin-mediated FASN inhibition and cancer cell suppression in MPM remains unclear.

This study aimed to determine promising diagnostic and therapeutic targets for *NF2/p16*-deficient MPM by FASN and its effect on mitochondrial dynamics. Additionally, considering that cerulenin exhibits a selective antiproliferative effect on MPM cells, its role as potential molecular-targeted anticancer drug was determined.

## Results

### The FASN inhibitor cerulenin selectively suppresses the proliferation of NF2-p16-deficient MPM cells

Given that we have previously shown differential gene expression between control and *NF2/p16*-double knockout (DKO) isogenic human mesothelial cell clones, we attempted to identify an inhibitor that specifically suppresses the proliferation of *NF2/p16*-DKO cells, which are frequently observed in MPM (38). To this end, we performed the MTT assay with the Screening Committee of the Anticancer Drugs library, which comprises 364 chemical compounds (Supplementary Table S1). Our drug screening assay showed that the FASN inhibitor cerulenin, but not C75, exhibited the most potent antiproliferative activity against *NF2/p16*-DKO cells (Fig. 1a). To further evaluate cerulenin’s impact on MPM cell proliferation, we conducted MTT assays using various mesothelial and MPM cell lines. These included *NF2*^+/+^/*p16*^+/+^ (parental MeT-5A and HOMC-B1 cells), *NF2*^−/−^/*p16*^+/+^ (Y-MESO-14), *NF2*^+/+^/*p16*^−/−^ (NCI-2452, ACC-MESO-4, Y-MESO-9, and MSTO-211H), and *NF2*^−/−^/*p16*^−/−^ cells (MeT-5A-DKO, HOMC-B1-DKO, NCI-H290, NCI-H2052, and MSTO-211H-DKO). Interestingly, the average IC_50_ values in the *NF2*/*p16*-DKO mesothelial cell lines and MSTO-211H cells ranged from 5 μM to 7.5 μM, which is lower than that in the *NF2*^+/+^*p16*^+/+^cell lines (approximately 20 μM) (Fig. 1b). This outcome indicates that cerulenin selectively inhibits the proliferation of *NF2*/*p16*-DKO cells. Compared with the antiproliferative effect of cerulenin, treatment with pemetrexed or cisplatin was relatively ineffective even at higher doses against both the *NF2*^+/+^*p16*^+/+^ and *NF2*^−/−^/*p16*^−/−^ MPM cell lines (Fig. 1c). Furthermore, cerulenin treatment suppressed the proliferation of MPM cell lines NCI-H290 and NCI-H2052, both of which spontaneously lacked *NF2*/*p16* expression (Fig. 1d). These findings indicate that cerulenin holds potential as a valuable molecular target for anticancer therapy in MPM.

**Fig. 1.**
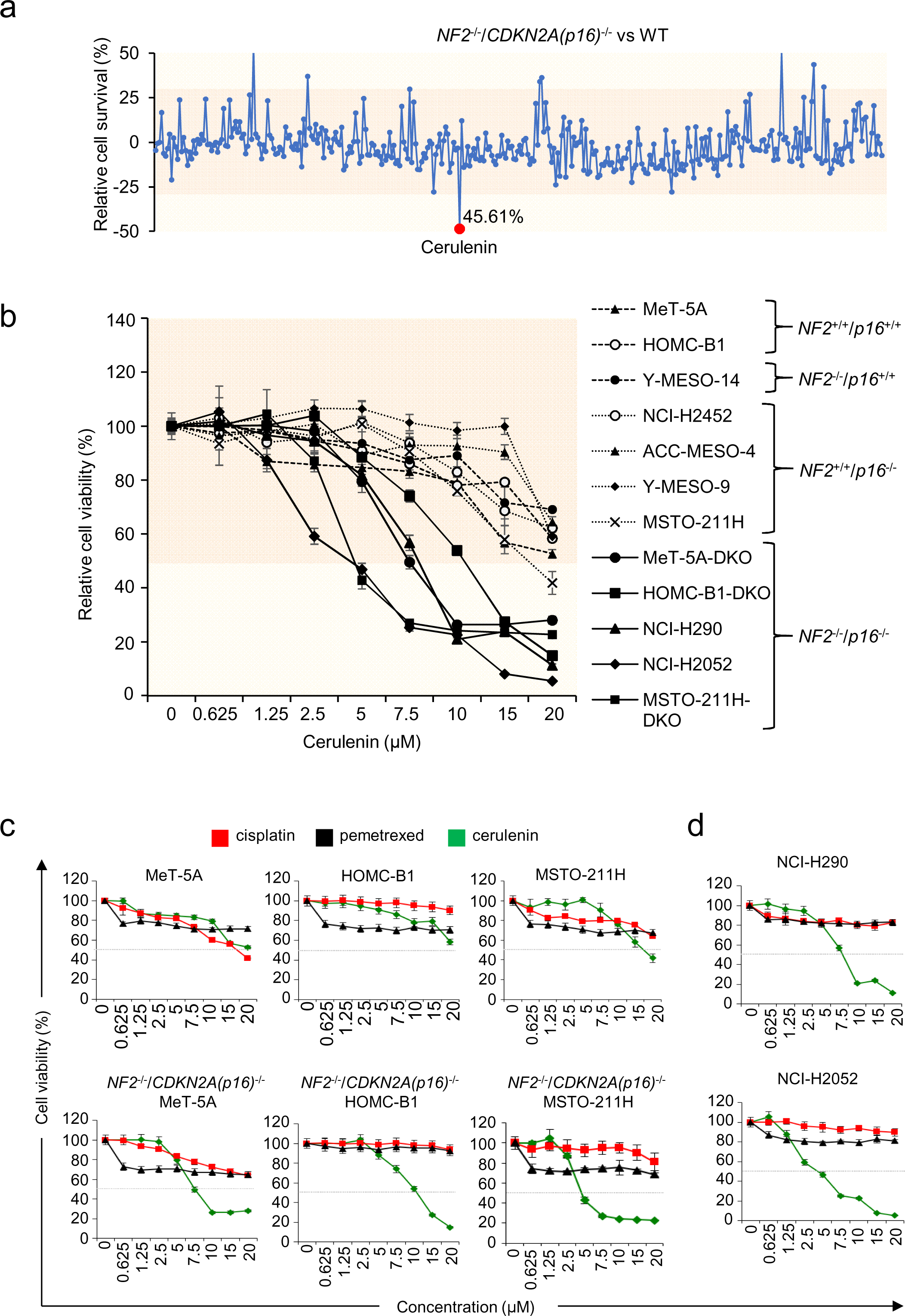
Identification of FASN inhibitor cerulenin against the DKO-deficient cells and its effect on MPM cell line survival. (**a**) Evaluation of compounds from the Screening Committee of Anticancer Drugs library. MeT-5A cells (DKO and NF2/p16-WT [parental] cells) were exposed to 364 chemical compounds (10 μM each). Cell survival rates were normalized to the mean optical densities of untreated cells (set as 100%). Results are depicted as differences in cell survival percentages between NF2/p16-WT and DKO cells. Red spots indicate >40% reduction in cell viability in DKO cells compared to NF2/p16-WT cells. (**b**) Assessment of the impact of the FASN inhibitor cerulenin on MPM cell line survival. Cell survival rates were calculated as described previously. (**c**) Examination of cisplatin, pemetrexed, and cerulenin effects on the survival of various MPM cell lines (MeT-5A, HOMC-B1, Y-MESO-14, Y-MESO-12, NCI-H2452, ACC-MESO-4, Y-MESO-9, MSTO-211H, MeT-5A-DKO, HOMC-B1-DKO, NCI-H290, NCI-H2052, and MSTO-211H-DKO) after incubation with indicated drug concentrations (20, 15, 10, 7.5, 5, 2.5, 1.25, 0.625, and 0 μM) for 72 hours. MTT assays were conducted following the manufacturer’s guidelines, and absorbance was measured at 595 nm using a spectrophotometer. Cell survival percentages were calculated accordingly. Cisplatin, pemetrexed, and cerulenin are denoted by red, black, and green lines, respectively. Data are presented as mean ± standard error (n = 3).

### FASN expression is inversely associated with NF2/p16 expression in human MPM tissues

Immunohistochemistry was performed using 45 MPM tissue samples and 2 mesothelium samples to assess FASN expression in mesothelioma (Supplementary Table S2). Microscopic analyses revealed 3 strong (3^+^), 14 moderate (2^+^), and 11 weak (1^+^) FASN-positive signals in the MPM tissues (Fig. 2a; Supplementary Table S2), whereas no positive signal was observed in the normal mesothelium (Fig. 2b). Additionally, the FASN positivity rate was higher in the NF2/p16-negative cases (15/15 tissues, 100%) than in the NF2/p16-positive cases (8/25 tissues, 32%; Fig. 2b). Notably, MPM patients demonstrating elevated FASN expression levels experienced notably shorter overall survival compared to those with lower FASN expression levels (Fig. 2c). These findings strongly imply a close correlation between FASN expression and poorer prognosis in MPM patients.

**Fig. 2.**
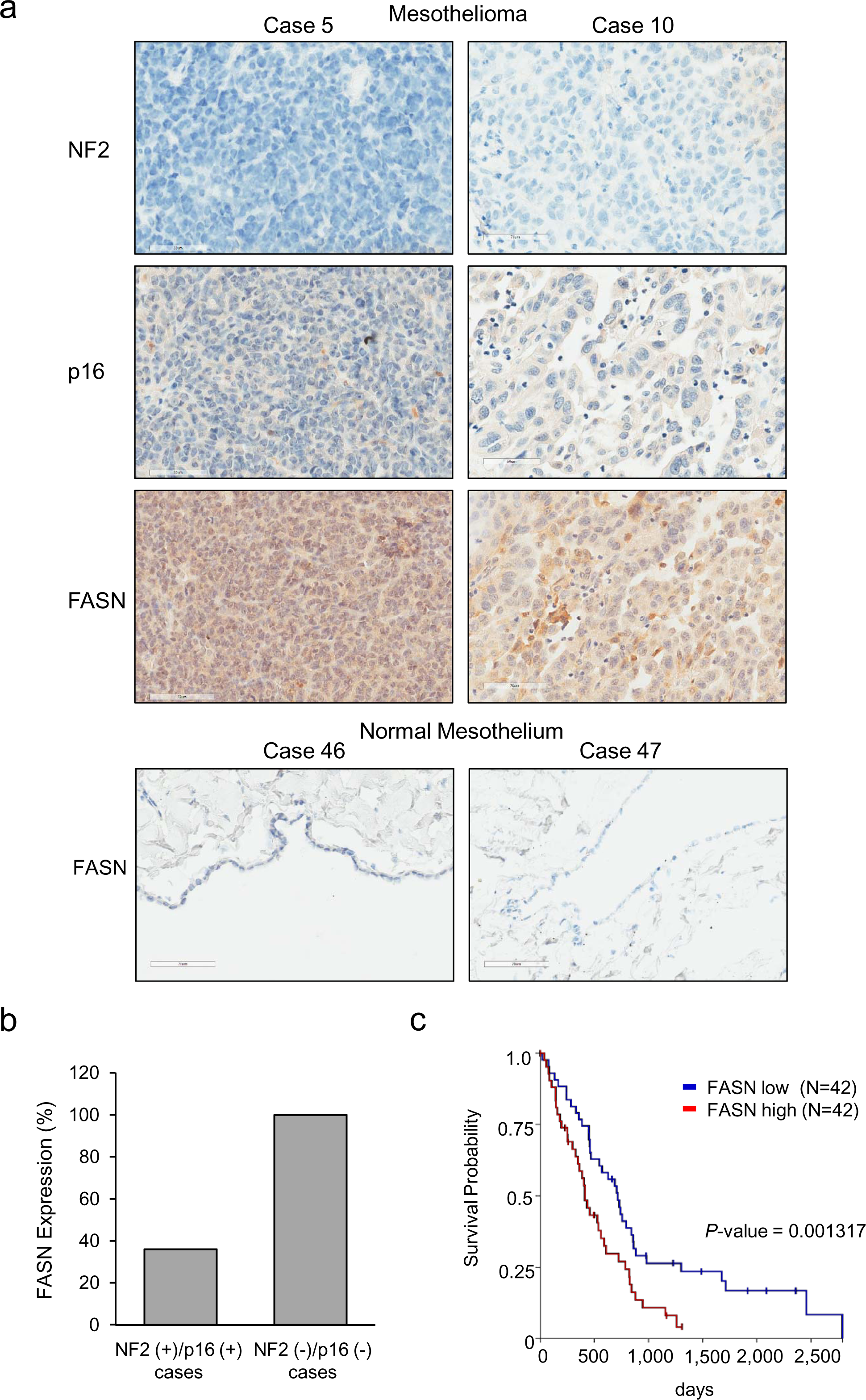
Immunohistochemical (IHC) analysis of FASN expression. (**a**) Illustrative IHC findings demonstrating FASN expression in two MPM tissues (top panels, cases 5 and 10) and two normal mesothelial tissues (bottom panel, cases 46 and 47). (**b**) Summary of IHC outcomes in MPM tissues. The bar graph indicates the percentage of total MPM cases exhibiting FASN expression by NF2/p16-positive or NF2/p16-negative cases. (**c**) Kaplan-Meier analysis was performed to assess the impact of FASN expression on overall survival in TCGA mesothelioma patients, sourced from the UCSC Xena database. Fluorescence values above and below the median were defined as high (red) and low (blue) FASN expression, respectively. Scale bar = 60 μm.

### Cerulenin suppresses tumor growth of NF/p16-deficient MSTO-211H cells in vivo

The effect on the growth of NF2/p16-deficient MSTO-211H (NF/p16-DKO MSTO-211H) tumors was examined in SCID mice to assess the potential of cerulenin for clinical application. The intraperitoneal injection of cerulenin notably inhibited tumor growth in comparison to the vehicle control (see Figs. 3a and 3b). Furthermore, cerulenin treatment did not result in any loss of body weight (see Fig. 3c). Subsequent assessment of cerulenin’s safety in normal BALB/c mice revealed no noteworthy alterations in body weight (data not presented) or histopathological changes in the heart, liver, or kidneys on Day 14 post-treatment (see Fig. 3d). Furthermore, biochemical analyses showed no alterations in blood parameters, including levels of aspartate aminotransferase (AST) and alanine aminotransferase (ALT), following cerulenin administration (see Fig. 3e; Supplementary Table S3). These findings indicate the potential safety and efficacy of cerulenin as a molecular-targeted anticancer medication for managing patients with NF2/p16-deficient MPM.

**Fig. 3.**
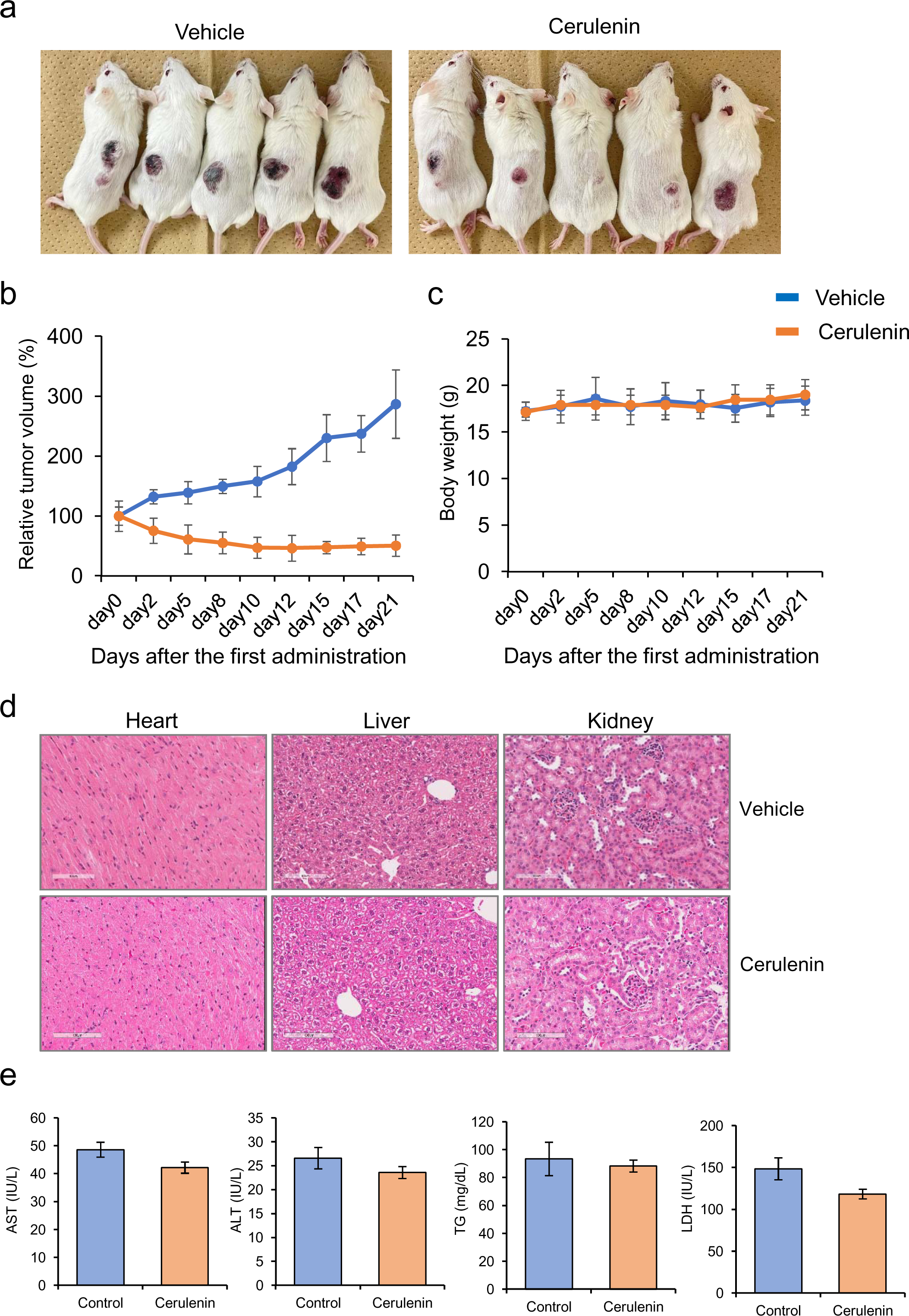
Effect of cerulenin on the growth of MSTO-211H-DKO tumor cells in vivo. The MSTO-211H-DKO cells (5 × 10^6^ cells/mouse) were injected subcutaneously into SCID mice. Following tumor establishment (day 0), cerulenin (20 mg/kg body weight) or PBS vehicle was administered intraperitoneally on days 0, 2, 5, 8, 10, 12, 15, and 17. (**a**) A representative image depicting tumor-bearing xenografted mice in each group. (**b–c**) Line graphs illustrating (**b**) relative tumor volume and (**c**) mouse body weight (in grams) during cerulenin treatment. Tumor volume is presented relative to the size at day 0 (defined as 100%). Data are presented as mean ± SE (n = 5). *p < 0.05. (**d**) Representative histochemical images of the heart, liver, and kidney from BALB/c mice after the 14-day observation period (Hematoxylin and Eosin staining, magnification, ×100; scale bar = 100 μm). (**e**) Alterations in blood chemistry (AST, ALT, TG, and LDH) following cerulenin administration compared with control data.

### Cerulenin accelerates the mitochondrial fusion processes and induces apoptosis in NF2/p16-DKO cells

Growing evidence indicates FASN’s role in tumor proliferation through the regulation of energy metabolism. Consequently, we investigated cerulenin’s impact on mitochondrial morphology in *NF2/p16*-DKO cells. We assessed the morphology of the mitochondria using MitoTracker, which permanently binds to mitochondria regardless of cell survival. Confocal microscopy showed that mitochondrial hyperfusion (long tubular network structure) increased after cerulenin treatment in *NF2/p16*-DKO MeT-5A and HOMC-B1 mesothelial cells, whereas no obvious change was observed in the parental cells (Fig. 4a). Similarly, mitochondrial hyperfusion increased after cerulenin treatment in parental MSTO-211H (spontaneously *CDKN2A/p16*-deficient) cells, which was further enhanced in the *NF2/p16*-DKO MSTO-211H cells (Fig. 4a). We examined whether cerulenin influences the phosphorylation status of dynamin-related protein 1 (DRP1), a critical regulator of mitochondrial fission, in *NF2/p16*-DKO cells. Immunofluorescence analysis revealed elevated levels of DRP1 phosphorylation in all *NF2/p16*-DKO cells compared to parental cells; however, cerulenin treatment mitigated these increases in DRP1 phosphorylation levels (see Fig. 4a). Additionally, Western blot analysis demonstrated that cerulenin reduced the phosphorylation levels of both Akt and DRP1 (see Fig. 4b). We unexpectedly found that the FASN protein levels were relatively higher in all the*NF2/p16*-DKO cells than in the parental cells tested (Fig. 4b). In fact, *NF2*/*p16* loss exhibited high FASN expression levels in the MPM cells tested, as compared with NF2- and/or p16-positive MPM cells (Supplementary Fig. S1). To assess cerulenin’s impact on the apoptosis of NF2/p16-double knockout (DKO) cells, we conducted flow cytometry analysis using Annexin V (AxV) and propidium iodide (PI) double staining. The results revealed a significant increase in the Ax^+^/PI^+^ population in NF2/p16-DKO mesothelial and MPM cells following cerulenin treatment compared to parental cells (see Fig. 4c), suggesting a potential heightened sensitivity of NF2/p16-DKO cells to cerulenin. Similarly, western blot analysis revealed that the cleaved poly (ADP-ribose) polymerase (PARP) levels markedly increased, whereas both FASN protein and phosphor-AKT levels decreased after cerulenin treatment in *NF2/p16*-DKO cells (Fig. 4b). Furthermore, our gene set enrichment analysis (GSEA) demonstrated a significant suppression of PI3K-AKT-MTOR signaling in NF2/p16-DKO cells treated with cerulenin (see Fig. 4d). These findings strongly indicate that cerulenin selectively triggers apoptosis by targeting oncogenic AKT signaling in NF2/p16-DKO cells.

**Fig. 4.**
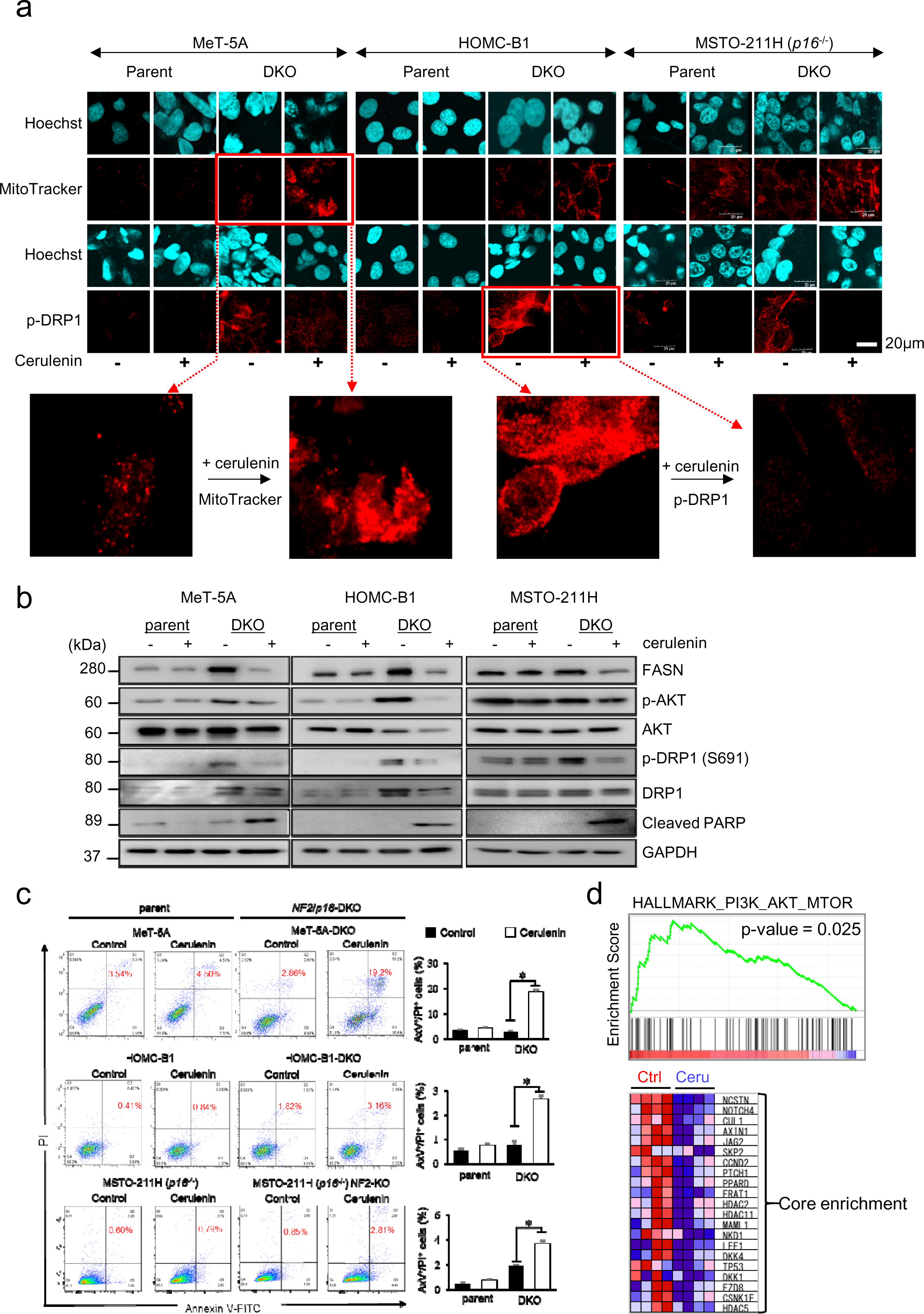
Role of cerulenin in mitochondrial morphology and apoptosis in DKO-deficient MPM cells. (**a**) The mitochondria were visualized using the MitoTracker probe (red). Nuclei were stained with Hoechst (blue), in MeT-5A (upper left panel), HOMC-B1 (upper center panel), and MSTO-211H (upper right panel). Cells were immunostained with Drp1 antibody (red) and Hoechst (blue) in the mitochondria, as depicted in the lower left panels (MeT-5A), lower center panels (HOMC-B1), and lower right panels (MSTO-211H). Two representative magnified view has been illustrated in bottom part. (**b**) Protein expression of FASN, p-AKT, AKT, p-Drp1, Drp1, c-PARP, and GAPDH analyzed by Western blotting in MeT-5A, MeT-5A-DKO, HOMC-B1, HOMC-B1-DKO, MSTO-211H, and MSTO-211H-DKO cells treated with cerulenin (7.5 µM) for 48 h. (**c**) Flow cytometry analysis. Representative results of AxV-PI-based staining are presented on the left. The graphs on the right display the percentage of AxV^+^/PI^+^ apoptotic cells following cerulenin treatment (7.5 µM) for 48 h measured using LSRFortessa X-20 Flow Cytometer (BD Biosciences). Data are mean ± SE (n = 3). Asterisks denote significant differences between the DKO-deficient cells (MeT-5A and HOMC-B1 cells and MSTO-211H cells) (*p < 0.05). (**d**) Gene set enrichment analysis (GSEA). MeT-5A, MeT-5A-DKO, HOMC-B1, and HOMC-B1-DKO cell clones were treated with cerulenin (7.5 µM) or control solvent for 48 h. A cDNA microarray analysis (Agilent) was conducted to obtain a comprehensive gene expression profile. Raw data were analyzed using GSEA with HALLMARK gene sets (C1).

### FASN knockout (TKO) suppresses cell proliferation and DRP1 activity in NF2/p16-DKO MeT-5A cells

*FASN* knockout (hereafter called triple knockout “TKO”) cell clones #1 and #2 were generated using previously established *NF2/p16*-DKO MeT-5A cell clones by targeting exon 3 of the *FASN* gene to further clarify the role of FASN in *NF2/p16*-DKO mesothelial cells (Supplementary Figs. 2a and b). As expected, the FASN protein expression levels were undetected in the TKO #1 and #2 cells (Supplementary Fig. S2c). Further, effect on the proliferation rate by the disruption of the *FASN* gene was tested. The MTT assay indicated a notably reduced proliferation rate in the TKO cell clones (#1 and #2) compared to the DKO cell clones (see Fig. 5a). Furthermore, Western blot analysis revealed a decrease in Akt and DRP1 phosphorylation levels, alongside elevated levels of cleaved PARP and caspase-3 in the TKO cell clones relative to the DKO cell clones (see Fig. 5b). Additionally, OPA1, MFN1, and MFN2, which positively regulate mitochondrial fusion, increased at the protein levels in the TKO cell clones (Fig. 5b). Therefore, the effect of *FASN* knockout on the mitochondrial morphology under *NF2/p16*-deficient conditions was examined. Confocal microscopy showed a significant increase in the elongated and tubular structure of the mitochondria in the TKO cells as compared with the DKO cells (Fig. 5c). The protein expression of DRP1 decreased consistently, whereas that of OPA1 increased in the TKO cell clones (Fig. 5d). The effect of proteasome inhibition on DRP1 protein expression was examined to uncover the molecular mechanism underlying the DRP1 expression change in the TKO cells. Western blot analysis demonstrated that both the phosphorylation and protein expression levels of DRP1 were reinstated following treatment with the proteasome inhibitor MG-132 in the TKO cells (see Fig. 5e). Furthermore, the immunoprecipitation assay revealed a higher DRP1 ubiquitin level in the TKO cells than in the DKO cells (Fig. 5f). The intracellular reactive oxygen species (ROS), largely generated in the mitochondria, increased in the TKO cells (Fig. 5g), suggesting involvement of FASN expression in the mitochondrial activity, which might affect mitochondrial ROS generation in *NF2/p16*-DKO cells.

**Fig. 5.**
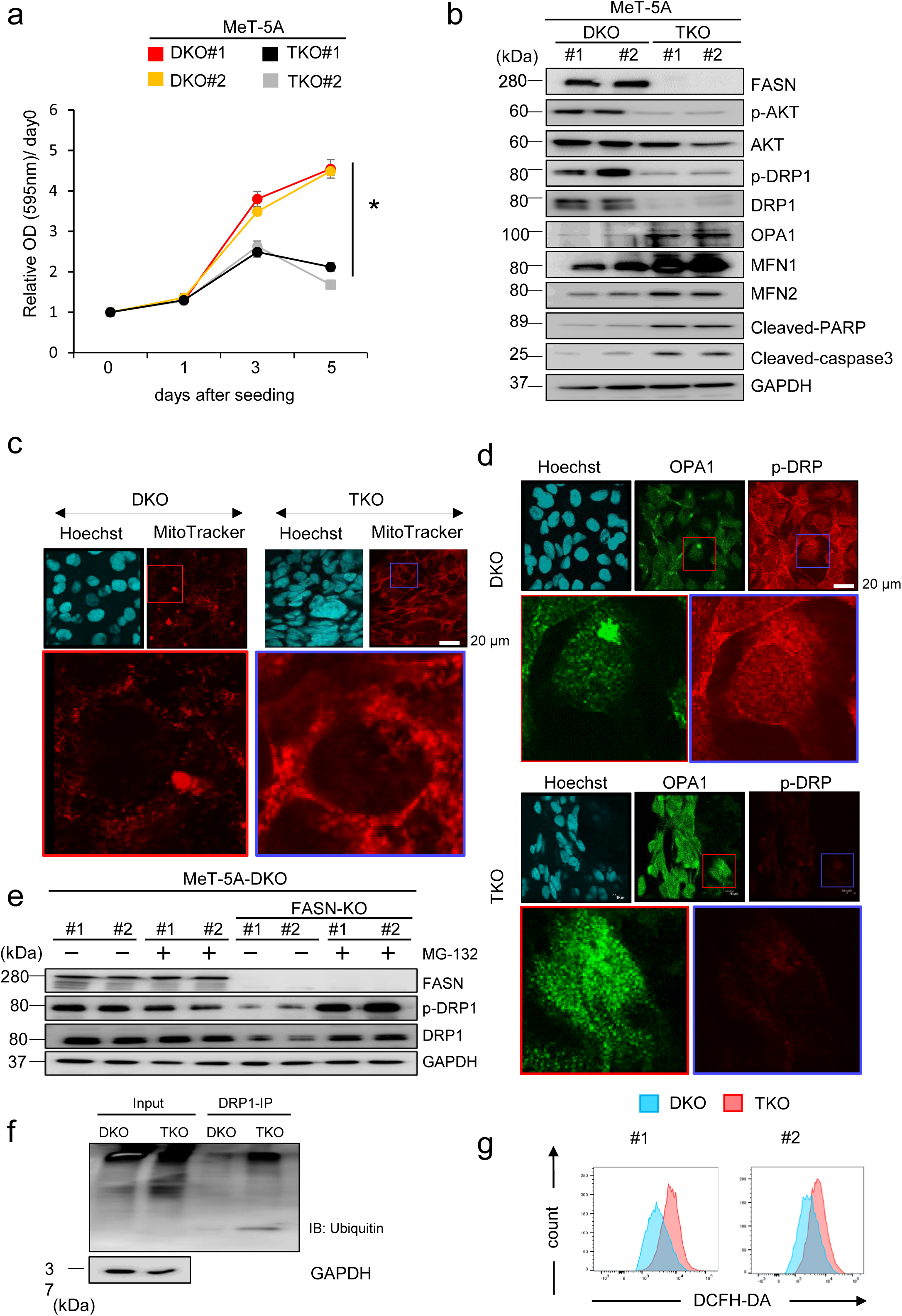
Disruption of *FASN* has adverse effects on mitochondrial morphology and dynamics in MPM cells. Cellular phenotype of MeT-5A-DKO/fatty acid synthase (DKO/FASN) triple knockout (TKO) in MeT-5A cells. (**a**) MTT assay of DKO (#1 and #2) and TKO (#1 and #2) cell clones in MeT-5A cells. The optical density (OD; 595 nm) at each time point (days 0, 1, 3, and 5) is presented as the mean ± SEM (n = 6). Growth ratios are expressed relative to the optical densities detected at day 0, arbitrarily defined as 1. (**b**) Western blot analysis of FASN, p-AKT, AKT, p-Drp1, Drp1, Opa1, MFN1, MFN2, c-PARP, and c-caspase3 expression in the DKO and TKO cells. (**c**) Mitochondria labeled with the MitoTracker probe (red; lower panel). Nuclei labeled with Hoechst (blue; upper panel). DKO and TKO shown in the left and right panels, respectively. (**d**) Cells immunostained with Opa1 antibody (green) and Hoechst (blue) in the mitochondria as shown in the left (DKO) and right (TKO) panels. (**e**) Effects of the proteasome inhibitor MG-132 on FASN, pDrp1, and Drp1 proteins observed in TKO and DKO cells. (**f**) DKO and TKO cells transfected with ubiquitin expression plasmid followed by MG-132 (10 μM) treatment for 6 hours. Subsequently, each cell lysate immunoprecipitated with Drp1 and subjected to immunoblot with anti-ubiquitin antibody (P4D1). (**g**) Effect of DCFH-DH (10 μM, 45 min) on reactive oxygen species production in TKO cells (Red) compared to DKO cells (Blue) in clone #1 (left panel) and #2 (right panel).

## Discussion

This study investigated the involvement of FASN in MPM cell proliferation and revealed the selective antiproliferative effect of the FASN inhibitor cerulenin in *NF2*/*p16*-deficient human MPM cells. The results revealed that loss of *NF2*/*p16* sensitizes the human mesothelial and MPM cells to cerulenin as compared with the *NF2*- and/or *p16*-intact cells.

Immunohistochemical analysis also showed detection of FASN expression in *NF2*/*p16*-negative human mesothelioma tissues. The analysis of public data revealed that the overall survival was significantly shorter for patients with MPM exhibiting a higher *FASN* expression level. Moreover, cerulenin suppressed the tumor growth of *NF2*/*p16*-deficient human MPM cells *in vivo* without any relevant toxicities.

FASN has been reported as a therapeutic target and an oncogene in several cancers, including breast, pancreatic, and colorectal cancers (38–43). FASN overexpression is correlated with poor prognosis in HER2-positive ovarian and gastric cancers (44, 45). FASN overexpression was observed in a subset of MPM cell lines and human MPM tissues in this study. Additionally, FASN expression was preferentially detected in the *NF2*/*p16*-deficient mesothelial and MPM cell lines as compared with the*NF2*/*p16*-intact mesothelial and MPM cell lines. MPM patients with higher FASN expression exhibited significantly shorter overall survival than those with lower FASN expression. Our previous study reported that the loss of *NF2* and *p16* genes significantly enhances anchorage-independent growth and epithelial– mesenchymal transition phenotype in mesothelial cells, in which the cancer-stem cell marker CD24 is upregulated in *NF2*/*p16*-deficient mesothelial and MPM cell lines (46, 13). Although the association between *NF2*/*p16* deficiency and FASN overexpression remains unclear, the retardation of the proliferation of *NF2*/*p16*-deficient cells by FASN disruption indicates the possibility of important role of FASN in *NF2*/*p16*-deficient MPM cell survival.

In our investigation, we observed that the upregulation of FASN expression coincides with the concurrent phosphorylation of oncogenic Akt in *NF2/p16*-DKO cells. Wagner et al. previously reported that the FASN inhibitor impairs receptor-PI3K-mTORC1 signaling in ovarian cancer cells (47). They also showed that FASN inhibitors impair the phosphatidylinositol 3,4,5-trisphosphate (PIP3) levels, diacylglycerol (DAG), and subsequent PI3K-AKT suppression (47). Another report showed that sensitivity to FASN inhibitors is driven by DAG level reductions and DAG-PKC signaling pathway impairments (48). We found that cerulenin significantly inactivates the PI3K–Akt–mTOR signaling pathway in *NF2*/*p16*-DKO mesothelial cells. Although it is still unclear whether the membrane lipid composition changes in the *NF2*/*p16*-DKO cells, the loss of *NF2*/*p16* may enhance the proliferation of MPM cells via FASN-mediated PI3K-Akt signaling. Indeed, *FASN* knockout in *NF2*/*p16*-DKO cells delays proliferation and decreases Akt phosphorylation as compared with the *FASN*-intact DKO cells. Therefore, it is possible that cerulenin suppresses PI3K-Akt signaling by modulating lipid membrane biogenesis.

The mitochondria are the main organs involved in the generation of cellular fuel ATP through several metabolic processes, including lipid catabolism. Besides, the equilibrium between mitochondrial fission and fusion occurs for the maintenance of mitochondrial functions, which also plays an important role in both normal cells and cancer progression (49–56). In this current study, the absence of FASN in the DKO (TKO) MeT-5A cells led to an increase in elongated and tubular structured mitochondria, indicative of mitochondrial fusion. Similarly, cerulenin treatment induces mitochondrial fusion formation in *NF2/p16*-deficient (DKO) MPM cells. The absence of FASN, as well as FASN inhibition, notably elevated the count of apoptotic cells in DKO MPM cells. These findings imply that FASN plays a crucial role in sustaining the survival of MPM cells through its involvement in mitochondrial activity. At a molecular level, both the expression and phosphorylation of DRP1 decreased in TKO MeT-5A cells and cerulenin-treated DKO cells. Furthermore, the depletion of FASN in TKO cells resulted in increased protein expression of OPA1, MFN1, and MFN2, all of which promote mitochondrial fusion. Interestingly, proteasome inhibitor MG-132 restored DRP1 protein expression in the TKO cells. Since the ubiquitinated DRP1 level is elevated in TKO cells compared to DKO cells, it suggests a potential role of FASN in stabilizing DRP1 at the protein level. Given that FASN-mediated fatty acids are converted to phospholipids (e.g., cardiolipin) that comprise the mitochondrial membrane, it would be of interest to examine whether cerulenin affects the lipid composition of mitochondrial membranes in MPM.

The present study has some limitations. First, the FASN protein expression was shown more frequent in NF2- and/or p16-negative MPM tissues as compared in both positive tissues; however, the somatic mutations of *NF2* and/or *p16* genes in the corresponding tissues was not investigated. Therefore, further studies are required to clarify the association between FASN expression and functional status of *NF2* and/or *p16* in MPM. Second, the antitumor effect of cerulenin was demonstrated using only one human MPM cell line, MSTO-211H. Hence, additional tests using primary human MPM-derived tumors are required for evaluating the efficacy of targeting FASN therapy in MPM.

In conclusion, our genome-edited human mesothelial cell model identified that the FASN inhibitor cerulenin preferentially exhibits an antiproliferative effect on MPM cells lacking NF2 and/or p16. These results underscore the potential clinical utility of cerulenin in crafting molecular-targeted therapies for combating MPM.

## Materials and methods

### Cell culture

Two immortalized normal human mesothelial cell lines, MeT-5A (pleural mesothelial) and HOMC-B1 (omental mesothelial; epithelioid type), along with eight human mesothelioma cell lines—ACC-MESO-4, Y-MESO-12, Y-MESO-14, Y-MESO-9, NCI-H2452, MSTO-211H, NCI-H290, and NCI-2052—were generously provided by Dr. Y. Sekido from the Division of Molecular Oncology, Aichi Cancer Center Research Institute (Nagoya, Japan). The HOMC-B1 cell lines were maintained 18s according to previously described protocols. The ACC-MESO-4, Y-MESO-12, Y-MESO-14, Y-MESO-9, NCI-H2452, MSTO-211H, NCI-H290, and NCI-2052 cell lines were cultured in RPMI-1640 medium (Wako Pure Chemical Industries, Ltd., Osaka, Japan) supplemented with 10% fetal bovine serum (Sigma) and 1% penicillin-streptomycin (Wako Pure Chemical Industries, Ltd.) at 37°C in a 5% CO2 humidified atmosphere.

### Gene KO using the CRISPR/Cas9 system

We employed the CRISPR/Cas9 system to disrupt NF2 expression in the MSTO-211H cell line and FASN expression in the DKO cells, following established procedures (12). The pSpCas9(BB)-2A-GFP (PX458) plasmid was generously provided by Feng Zhang (plasmid #48138; Addgene, Watertown, MA, USA) (58). Briefly, sgRNA sequences were chosen using an optimized CRISPR design tool (http://crispr.mit.edu/). The selected sgRNA sequences for NF2 and FASN were 5′-AAACATCTCGTACAGTGACA-3′ in exon 8 and 5′-CCTTCAGCTTGCCGGACCGC-3′ in exon 3, respectively. Plasmids expressing hCas9 and sgRNA were generated by ligating oligonucleotides into the BbsI site of PX458 (NF2/PX458 or FASN/PX458). To establish a knockout (KO) clone, 1 μg of NF2/PX458 plasmid was transfected into the MSTO-211H cell line, and 1 μg of FASN/PX458 plasmid was transfected into MeT-5A-DKO#1 cells (1 × 10^6^ cells) using a 4D-Nucleofector instrument (Lonza Japan, Tokyo, Japan). After 3 days, GFP-expressing cells were sorted using fluorescence-activated cell sorting (BD FACSAria™ III Cell Sorter; BD Biosciences, San Jose, CA, USA). A single clone was selected, expanded, and utilized for the biological assays.

### Cell growth assay

The growth rate of the cells was assessed using the MTT assay (59). Briefly, cells (1 × 10^3 cells/well) were seeded into 96-well plates and cultured for the specified durations. Subsequently, 10 μL of MTT solution (5 mg/mL; Sigma-Aldrich) was added to each well, and the cells were incubated for 4 hours. Following this, cell lysis buffer was added to dissolve the resultant colored formazan crystals. The absorbance at 595 nm was measured using a SpectraMAX M5 spectrophotometer (Molecular Devices, Sunnyvale, CA, USA).

### Western blot analysis

Western blotting was conducted following established protocols (13, 60). The antibodies utilized are detailed in Table S4. Immune complexes were visualized using ImmunoStar LD (Wako Pure Chemical Industries, Ltd.) and imaged with a LAS-4000 image analyzer (GE Healthcare, Tokyo, Japan).

### Annexin V assay

Cells were plated into six-well culture plates (5 × 10^5 cells/well) and treated with cerulenin (7.5 µg/mL) for 48 hours. Subsequently, the cells were exposed to annexin V (Ax)-FITC and propidium iodide (PI) (10 μg/mL) at 25°C for 15 minutes. The fluorescence intensities were quantified using fluorescence-activated cell sorting (FACS) analysis (LSRFortessa X-20 Flow Cytometer, BD Biosciences,, Franklin Lakes, NJ, USA).

### Immunofluorescence

The cells were cultured on glass coverslips and fixed with a 4% paraformaldehyde solution for 20 minutes at 25°C. Subsequently, the cells were permeabilized with phosphate-buffered saline (PBS) containing 0.1% Triton X-100, blocked in PBS containing 7% serum for 30 minutes, and then incubated with primary antibodies followed by Alexa Fluor-conjugated secondary antibodies (Invitrogen). Cell staining was conducted using MitoTracker (stock solution 1 mM; diluted at 1:10,000) for 1 hour at 37°C to visualize the mitochondria. Following staining, the cells were washed with PBS and fixed with cold paraformaldehyde (3.2% in PBS) for 20 minutes at room temperature. After additional washing steps, the samples were mounted using PermaFluor, and images were captured using the FLUOVIEW FV3000 Series of Confocal Laser Scanning Microscopes.

### Immunohistochemistry

Immunohistochemical analysis was conducted following previously established protocols (61). A human mesothelioma tissue array (MS-1001a; US Biomax, Rockville, MD, USA) was utilized. The tissue sections were incubated with primary antibodies (NF2, p16, and FASN antibody, 2 μg/mL). Negative controls were established using normal rabbit immunoglobulin G or by omitting the primary antibody. Immunoreactivity was independently assessed by two investigators (S.K. and H.M.), and staining intensity was graded as strong (+3), moderate (+2), weak (+1), or negative (0).

### Xenograft experiments

All animal experiments were conducted in accordance with the protocols (approval number-2022-20) approved by the ethical committee of Aichi Medical University and followed established guidelines. Female Fox Chase SCID mice (CB17/Icr-Prkdcscid/IcrIcoCrl) were procured from CLEA Japan Inc. (Tokyo, Japan) and housed at the Institute of Animal Experiments in Aichi Medical University under pathogen-free conditions. The mice, aged 6–8 weeks and weighing 17–18 g each, were utilized for the study. Subcutaneous injection of MSTO-211H-DKO cells (6 × 10^6^ cells) was performed in the SCID mice. Upon reaching a tumor volume of approximately ∼100 mm^3^, the mice were randomly assigned to two groups (treatment and control). Cerulenin (20 mg/kg, administered three times per week) was intraperitoneally administered to mice in the treatment group, while PBS served as the vehicle control in the control group. Tumor measurements were taken every 3–4 days, and tumor volume was calculated using the modified ellipsoid formula (1/2 × length × width^2^).

### DCFH–DA-based DCF assay

Cells were plated into six-well culture plates at a density of 3 × 10^5^ cells per well. The following day, they were exposed to 10 μM DCFH–DA (2’,7’-dichlorodihydrofluorescein diacetate) in the medium for 45 minutes in the absence of light. Subsequently, the DCFH–DA solution was removed, and the cells were washed with PBS. After detachment using trypsin, the cells were suspended in media, centrifuged at 1,000 rpm for 5 minutes, and resuspended in PBS in 0.5-mL tubes, which were then placed on ice. The samples were analyzed via FACS using LSRFortessa X-20 Flow Cytometer (BD Biosciences).

### Statistical analysis

The data are presented as mean ± SE values. Statistical significance among groups was assessed using one-way analysis of variance followed by Dunnett’s comparison. All statistical analyses were conducted using SPSS 23.0 software (SPSS Inc., Chicago, Illinois, USA).

## Abbreviations

MPM: malignant pleural mesothelioma
IHC: immunohistochemistry
CRISPR: clustered regularly interspaced short palindromic repeats
Cas9: CRISPR-associated protein 9
FASN: Fatty acid synthase
DKO: double knockout
TKO: FASN-knockout DKO
MTR: MitoTracker
Drp1: dynamin-related protein 1
OPA1: optic atrophy-1
PARP: Poly (ADP-ribose) polymerase
ROS: reactive oxygen species
FCM: flow cytometry.

## Competing interest

This publication has no conflicts of interest, and there has been no substantial financial support for this work that could have influenced its outcome. The manuscript has been reviewed and endorsed by all named authors, and there are no other individuals who meet the criteria for authorship but are not listed. All authors have consented to the order in which they are listed in the manuscript.

## Availability of data and materials

All accessible data are provided either in the main manuscript or in the supplementary materials. Complete and unaltered western blot data utilized in the study are available in the supplementary information. Furthermore, specific inquiries, data, and materials can be obtained upon reasonable request to the corresponding author.

## Ethical standards

The research conducted adhered to the ethical standards outlined by the Japanese Ministry of Health, Labour, and Welfare.

## Author contributions

The study concept was devised by S.K. and A.O. oversaw project acquisition and supervision. S.K., M.L.R., M.N.H., and M.W. formulated the statistical analysis plan and performed the statistical analyses. Data and resources were acquired by S.K., H.M., L.Q.V., I.H., M.T.A.S., N.J., A.I., M.R., H.I., Y.K., Y.L., T.H., H.K., S.T., and Y.H. Acquisition of funding was facilitated by S.K. and A.O. contributed to manuscript composition. All authors critically reviewed and edited the manuscript draft and consented to the final version for publication.

We express our gratitude to Dr. Y. Sekido from the Division of Molecular Oncology, Aichi Cancer Center Research Institute, for generously providing the normal mesothelial cell lines (MeT-5A and HOMC-D4). This study received partial support from grants provided by the Ministry of Education, Culture, Sports, and Technology of Japan (MEXT, 19K08668, 22K08294 to Y.H., 19K09292, 22K08985 to SK, and 21K08426 to AO), a research grant from the Hori Science and Arts Foundation, and a research grant from the Hirose International Scholarship Foundation (SK). MNH were supported by the Japanese Government (MEXT) Scholarship for Research Students.

## Notes

### Competing Interest Statement

The authors have declared no competing interest.

### Summary of Updates

Figure 5b, western blot image was not visible in PDF file and hence reuploaded the corrected version

## References

1. Carbone, M. et al. Malignant mesothelioma: facts, myths, and hypotheses. J. Cell Physiol. 227, 44–58 (2012).

2. Roushdy-Hammady, I., Siegel, J., Emri, S., Testa, J. R., & Carbone M. Genetic-susceptibility factor and malignant mesothelioma in the Cappadocian region of Turkey. Lancet 357(9254), 444–5 (2001).

3. Carbone, C. et al. Mesothelioma: Scientific clues for prevention, diagnosis, and therapy. CA-Cancer J. Clin 69, 402–429 (2019).

4. Patel SC, Dowell JE. Modern management of malignant pleural mesothelioma. Lung. Sci. U S A. 113, 13432–13437 (2016).

5. Vogelzang NJ, Rusthoven JJ, Symanowski J, et al. Phase III study of pemetrexed in combination with cisplatin versus cisplatin alone in patients with malignant pleural mesothelioma. J Clin Oncol.

6. Fung, H., Kow, Y. W., Van, H. B. & Mossman, B. T. Patterns of 8 hydroxy deoxyguanosine formation in DNA and indications of oxidative stress in rat and human pleural mesothelial cells after exposure to crocidolite asbestos. Carcinogenesis 18, 825–832 (1997).

7. Sekido, Y. Molecular pathogenesis of malignant mesothelioma. Carcinogenesis 34, 1413– 1419 (2013).

8. Testa, J. R. et al. Germline BAP1 mutations predispose to malignant mesothelioma. Nat. Genet. 43, 1022–1025 (2011).

9. Bott, M. et al. The nuclear deubiquitinase BAP1 is commonly inactivated by somatic mutations and 3p21.1 loss in malignant pleural mesothelioma. Nat. Genet. 43, 668–672 (2011).

10. Guo, G. et al. Whole-exome sequencing reveals frequent genetic alterations in BAP1, NF2, CDKN2A, and CUL1 in malignant pleural mesothelioma. Cancer Res. 75, 264–269 (2015).

11. Bianchi, A. B., et al. High frequency of inactivating mutations in the neurofibromatosis type 2 gene (NF2) in primary malignant mesotheliomas. Proc. Natl Acad. Sci. USA 92, 10854–10858 (1995).

12. Wahiduzzaman, M. et al. Establishment and characterization of CRISPR/Cas9-mediated NF2–/– human mesothelial cell line: Molecular insight into fibroblast growth factor receptor 2 in malignant pleural mesothelioma. Cancer Sci. 110, 180–193. (2019).

13. Karnan, S., Ota, A., Murakami, H. et al. Identification of CD24 as a potential diagnostic and therapeutic target for malignant pleural mesothelioma. Cell Death Discov. 6, 127 (2020).

14. Karnan, S., Ota, A., Murakami H, et al. CAMK2D: a novel molecular target for BAP1-deficient malignant mesothelioma. Cell Death Discov. 9, 257 (2023).

15. Menendez JA, Lupu R. Fatty acid synthase and the lipogenic phenotype in cancer pathogenesis. Nat. Rev. Cancer. 2007;7(10):763–777.

16. Kuhajda FP. Fatty-acid synthase and human cancer: new perspectives on its role in tumor biology. Nutrition. 2000;16(3):202–208.

17. Ho TS, Ho YP, Wong WY, Chi-Ming Chiu L, Wong YS, Eng-Choon Ooi V. Fatty acid synthase inhibitors cerulenin and C75 retard growth and induce caspase-dependent apoptosis in human melanoma A-375 cells. Biomed Pharmacother. 2007.

18. Schroeder B, Vander Steen T, Espinoza I, Venkatapoorna CMK, Hu Z, Silva FM, Regan K, Cuyàs E, Meng XW, Verdura S, Arbusà A, Schneider PA, Flatten KS, Kemble G, Montero J, Kaufmann SH, Menendez JA, Lupu R. Fatty acid synthase (FASN) regulates the mitochondrial priming of cancer cells. Cell Death Dis. 2021 Oct 21;12(11):977.

19. Fafián-Labora J, Carpintero-Fernández P, Jordan SJD, Shikh-Bahaei T, Abdullah SM, Mahenthiran M, Rodríguez-Navarro JA, Niklison-Chirou MV, O’Loghlen A. FASN activity is important for the initial stages of the induction of senescence. Cell Death Dis. 2019 Apr 8;10(4):318.

20. Zaytseva YY, Harris JW, Mitov MI, Kim JT, Butterfield DA, Lee EY, Weiss HL, Gao T, Evers BM. Increased expression of fatty acid synthase provides a survival advantage to colorectal cancer cells via upregulation of cellular respiration. Oncotarget. 2015 Aug 7;6(22):18891–904.

21. Wu D, Yang Y, Hou Y, Zhao Z, Liang N, Yuan P, Yang T, Xing J, Li J. Increased mitochondrial fission drives the reprogramming of fatty acid metabolism in hepatocellular carcinoma cells through suppression of Sirtuin 1. Cancer Commun (Lond). 2022 Jan;42(1):37–55.

22. Srinivasan S, Guha M, Kashina A & Avadhani NG Mitochondrial dysfunction and mitochondrial dynamics-The cancer connection. Biochim Biophys Acta Bioenerg 1858, 602–614 (2017).

23. Mannella CA. The relevance of mitochondrial membrane topology to mitochondrial function. Biochim Biophys Acta 1762, 140–147 (2006).

24. Johnson J, Mercado-Ayon E, Mercado-Ayon Y, Dong YN, Halawani S, Ngaba L et al. Mitochondrial dysfunction in the development and progression of neurodegenerative diseases. Arch Biochem Biophys 702, 108698 (2021).

25. Portz P & Lee MK. Changes in Drp1 function and mitochondrial morphology are associated with the _α_-synuclein pathology in a transgenic mouse model of Parkinson’s disease. Cells 10, 885 (2021).

26. Zhan X, Yu W, Franqui-Machin R, Bates ML, Nadiminti K, Cao H et al. Alteration of mitochondrial biogenesis promotes disease progression in multiple myeloma. Oncotarget 8, 111213–111224 (2017).

27. Ortiz-Ruiz A, Ruiz-Heredia Y, Morales ML, Aguilar-Garrido P, García-Ortiz A, Valeri A et al. Myc-related mitochondrial activity as a novel target for multiple myeloma. Cancers (Basel) 13 (2021).

28. Lv S, Zhang Y, Song J, Chen J, Huang B, Luo Y, Zhao Y. Cerulenin suppresses ErbB2-overexpressing breast cancer by targeting ErbB2/PKM2 pathway. Med Oncol. 2022 Oct 29;40(1):5. doi: 10.1007/s12032-022-01872-z. PMID: 36308575.

29. Wang Q, Du X, Zhou B, Li J, Lu W, Chen Q, Gao J. Mitochondrial dysfunction is responsible for fatty acid synthase inhibition-induced apoptosis in breast cancer cells by PdpaMn. Biomed Pharmacother. 2017 Dec;96:396–403. doi: 10.1016/j.biopha.2017.10.008. Epub 2017 Oct 12. PMID: 29031197.

30. Chang L, Wu P, Senthilkumar R, Tian X, Liu H, Shen X, Tao Z, Huang P. Loss of fatty acid synthase suppresses the malignant phenotype of colorectal cancer cells by down-regulating energy metabolism and mTOR signaling pathway. J Cancer Res Clin Oncol. 2016 Jan;142(1):59–72. doi: 10.1007/s00432-015-2000-8. Epub 2015 Jun 25. PMID: 26109148.

31. Murata S, Yanagisawa K, Fukunaga K, Oda T, Kobayashi A, Sasaki R, Ohkohchi N. Fatty acid synthase inhibitor cerulenin suppresses liver metastasis of colon cancer in mice. Cancer Sci. 2010 Aug;101(8):1861–5. doi: 10.1111/j.1349-7006.2010.01596.x. Epub 2010 Apr 21. PMID: 20491775.

32. Uddin S, Hussain AR, Ahmed M, Bu R, Ahmed SO, Ajarim D, Al-Dayel F, Bavi P, Al-Kuraya KS. Inhibition of fatty acid synthase suppresses c-Met receptor kinase and induces apoptosis in diffuse large B-cell lymphoma. Mol Cancer Ther. 2010 May;9(5):1244–55. doi: 10.1158/1535-7163.MCT-09-1061.

33. Deepa PR, Vandhana S, Krishnakumar S. Fatty acid synthase inhibition induces differential expression of genes involved in apoptosis and cell proliferation in ocular cancer cells. Nutr Cancer. 2013;65(2):311–6. doi: 10.1080/01635581.2013.748923. PMID: 23441619.

34. De Schrijver E, Brusselmans K, Heyns W, Verhoeven G and Swinnen JV: RNA interference-mediated silencing of the fatty acid synthase gene attenuates growth and induces morphological changes and apoptosis of LNCaP prostate cancer cells. Cancer Res 63: 3799–3804, 2003.

35. Gabrielson EW, Pinn ML, Testa JR and Kuhajda FP: Increased fatty acid synthase is a therapeutic target in mesothelioma. ClinCancer Res 7: 153–177, 2001.

36. Heiligtag, S., Bredehorst, R. & David, K. Key role of mitochondria in cerulenin-mediated apoptosis. Cell Death Differ 9, 1017–1025 (2002). 10.1038/sj.cdd.4401055

37. Jeong NY and Jeong NY: Cerulenin-induced apoptosis is mediated by disrupting the interaction between AIF and hexokinase II. Int J Oncol 40: 1949–1956, 2012.

38. Flavin R, Peluso S, Nguyen PL, Loda M. Fatty acid synthase as a potential therapeutic target in cancer. Future Oncol. 2010;6(4):551–62.

39. Menendez JA, Lupu R. Fatty acid synthase and the lipogenic phenotype in cancer pathogenesis. Nat Rev Cancer. 2007;7(10):763–77.

40. Alo, P. L., et al. Alo PL, Amini M, Piro F, Pizzuti L, Sebastiani V, Botti C, Murari R, Zotti G, Di Tondo U. Immunohistochemical expression and prognostic significance of fatty acid synthase in pancreatic carcinoma. Anticancer Res. 27(4B):2523–7. 2007.

41. Notarnicola M, Tutino V, Calvani M, Lorusso D, Guerra V, Caruso MG. Serum levels of fatty acid synthase in colorectal cancer patients are associated with tumor stage. J Gastrointest Cancer. 2012 Sep;43(3):508–11.

42. Wang Y, Kuhajda FP, Li JN, Pizer ES, Han WF, Sokoll LJ, Chan DW. Fatty acid synthase (FAS) expression in human breast cancer cell culture supernatants and in breast cancer patients. Cancer Lett. 167(1):99–104. 2001.

43. Kuhajda FP, Jenner K, Wood FD, Hennigar RA, Jacobs LB, Dick JD, Pasternack GR. Fatty acid synthesis: a potential selective target for antineoplastic therapy. Proc Natl Acad Sci U S A. 91(14):6379–83. 1994.

44. Cai Y, Wang J, Zhang L, Wu D, Yu D, Tian X, Liu J, Jiang X, Shen Y, Zhang L, Ren M, Huang P. Expressions of fatty acid synthase and HER2 are correlated with poor prognosis of ovarian cancer. Med Oncol. 32(1):391, 2015.

45. Castagnoli L, Corso S, Franceschini A, Raimondi A, Bellomo SE, Dugo M, Morano F, Prisciandaro M, Brich S, Belfiore A, Vingiani A, Di Bartolomeo M, Pruneri G, Tagliabue E, Giordano S, Pietrantonio F, Pupa SM. Fatty acid synthase as a new therapeutic target for HER2-positive gastric cancer. Cell Oncol (Dordr). 46(3):661–676, 2023.

46. Jaggupilli A, Elkord E. Significance of CD44 and CD24 as cancer stem cell markers: an enduring ambiguity. Clin Dev Immunol. 708036, 2012.

47. Wagner R, Stübiger G, Veigel D, Wuczkowski M, Lanzerstorfer P, Weghuber J, Karteris E, Nowikovsky K, Wilfinger-Lutz N, Singer CF, Colomer R, Benhamú B, López-Rodríguez ML, Valent P, Grunt TW. Multi-level suppression of receptor-PI3K-mTORC1 by fatty acid synthase inhibitors is crucial for their efficacy against ovarian cancer cells. Oncotarget. 8(7):11600–11613, 2017.

48. Benjamin DI, Li DS, Lowe W, Heuer T, Kemble G, Nomura DK. Diacylglycerol Metabolism and Signaling Is a Driving Force Underlying FASN Inhibitor Sensitivity in Cancer Cells. ACS Chem Biol. 10(7):1616–23, 2015.

49. Yaffe MP (1999) The machinery of mitochondrial inheritance and behavior. Science 283: 1493–1497

50. Shaw JM, Nunnari J (2002) Mitochondrial dynamics and division in budding yeast. Trends Cell Biol 12: 178–184

51. Chen H, Detmer SA, Ewald AJ, Griffin EE, Fraser SE, Chan DC (2003) Mitofusins Mfn1 and Mfn2 coordinately regulate mitochondrial fusion and are essential for embryonic development. J Cell Biol 160: 189–200

52. Misaka T, Miyashita T, Kubo Y (2002) Primary structure of a dynamin-related mouse mitochondrial GTPase and its distribution in brain, subcellular localization, and effect on mitochondrial morphology. J Biol Chem 277: 15834–15842

53. Smirnova E, Griparic L, Shurland DL, van der Bliek AM (2001) Dynamin-related protein Drp1 is required for mitochondrial division in mammalian cells. Mol Biol Cell 12: 2245–2256

54. James DI, Parone PA, Mattenberger Y, Martinou JC (2003) hFis1, a novel component of the mammalian mitochondrial fission machinery. J Biol Chem 278: 36373–36379

55. Boulton DP, Caino MC (2022) Mitochondrial Fission and Fusion in Tumor Progression to Metastasis. Front Cell Dev Biol. 10:849962

56. Karnan S, Hanamura I, Ota A, Vu LQ, Uchino K, Horio T, Murakami S, Mizuno S, Rahman ML, Wahiduzzaman M, Hasan MN, Biswas M, Hyodo T, Ito H, Suzuki A, Konishi H, Tsuzuki S, Hosokawa Y, Takami A. (2024) ARK5 enhances cell survival associated with mitochondrial morphological dynamics from fusion to fission in human multiple myeloma cells. Cell Death Discov 10:56

57. Sato T, Nakanishi H, Akao K, Okuda M, Mukai S, Kiyono T, Sekido Y. Three newly established immortalized mesothelial cell lines exhibit morphological phenotypes corresponding to malignant mesothelioma epithelioid, intermediate, and sarcomatoid types, respectively. Cancer Cell Int. 2021 Oct 18;21(1):546.

58. Ran FA, Hsu PD, Wright J, Agarwala V, Scott DA, Zhang F (2013) Genome engineering using the CRISPR-Cas9 system. Nat Protoc 8(11):2281–2308.

59. Hasan MN, Hyodo T, Biswas M, Rahman ML, Mihara Y, Karnan S, et al. Flow cytometry-based quantification of genome editing efficiency in human cell lines using the L1CAM gene. PLOS ONE. 2023;18(11):e0294146.

60. Karnan S, Hanamura I, Ota A, Takasugi S, Nakamura A, Takahashi M, Uchino K, Murakami S, Wahiduzzaman M, Quang Vu L, Rahman ML, Hasan MN, Hyodo T, Konishi H, Tsuzuki S, Yoshikawa K, Suzuki S, Ueda R, Ejiri M, Hosokawa Y, Takami A. (2021) CD52 is a novel target for the treatment of FLT3-ITD-mutated myeloid leukemia. Cell Death Discov 7(1):121

61. Ito T, Matsubara D, Tanaka I, Makiya K, Tanei ZI, Kumagai Y, Shiu SJ, Nakaoka HJ, Ishikawa S, Isagawa T, Morikawa T, Shinozaki-Ushiku A, Goto Y, Nakano T, Tsuchiya T, Tsubochi H, Komura D, Aburatani H, Dobashi Y, Nakajima J, Endo S, Fukayama M, Sekido Y, Niki T, Murakami Y (2016) Loss of YAP1 defines neuroendocrine differentiation of lung tumors. Cancer Sci 107(10):1527–1538

