## Supplementary material for "Targeting fatty acid synthase suppresses tumor development in *NF2/CDKN2A*-deficient malignant pleural mesothelioma": Supplemmental information (from another version of this preprint: version 2, 2024-07-18)

**Manuscript Title:**

- Supplementary information includes 6 supplemental figures and 4 supplemental table.

- All the full and uncropped version of the western blot images used in the manuscript has been included in this file (Supplemental Fig. S3~ Supplemental Fig. S6)

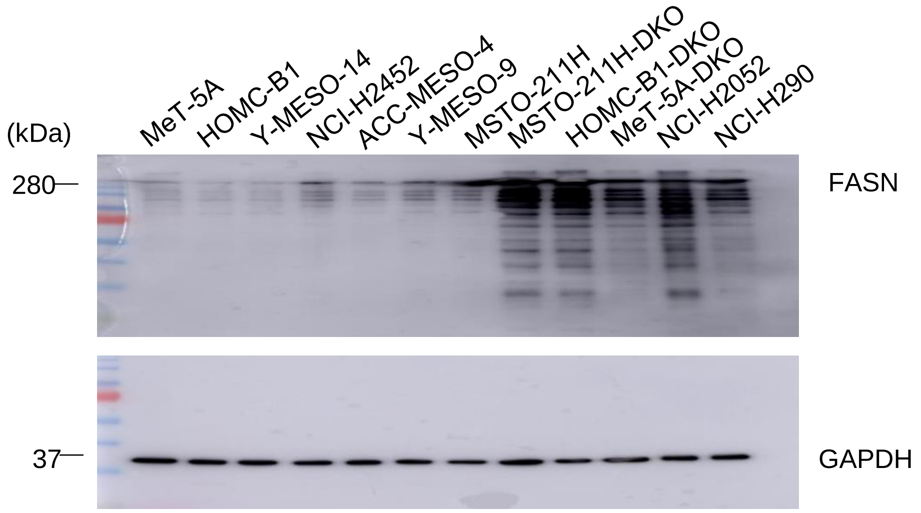


**Supplementary Fig. S1.** Western blot analysis was conducted to assess FASN expression in both mesothelial and MPM cell lines, with GAPDH serving as the internal control.

### Supplementary Fig. S2

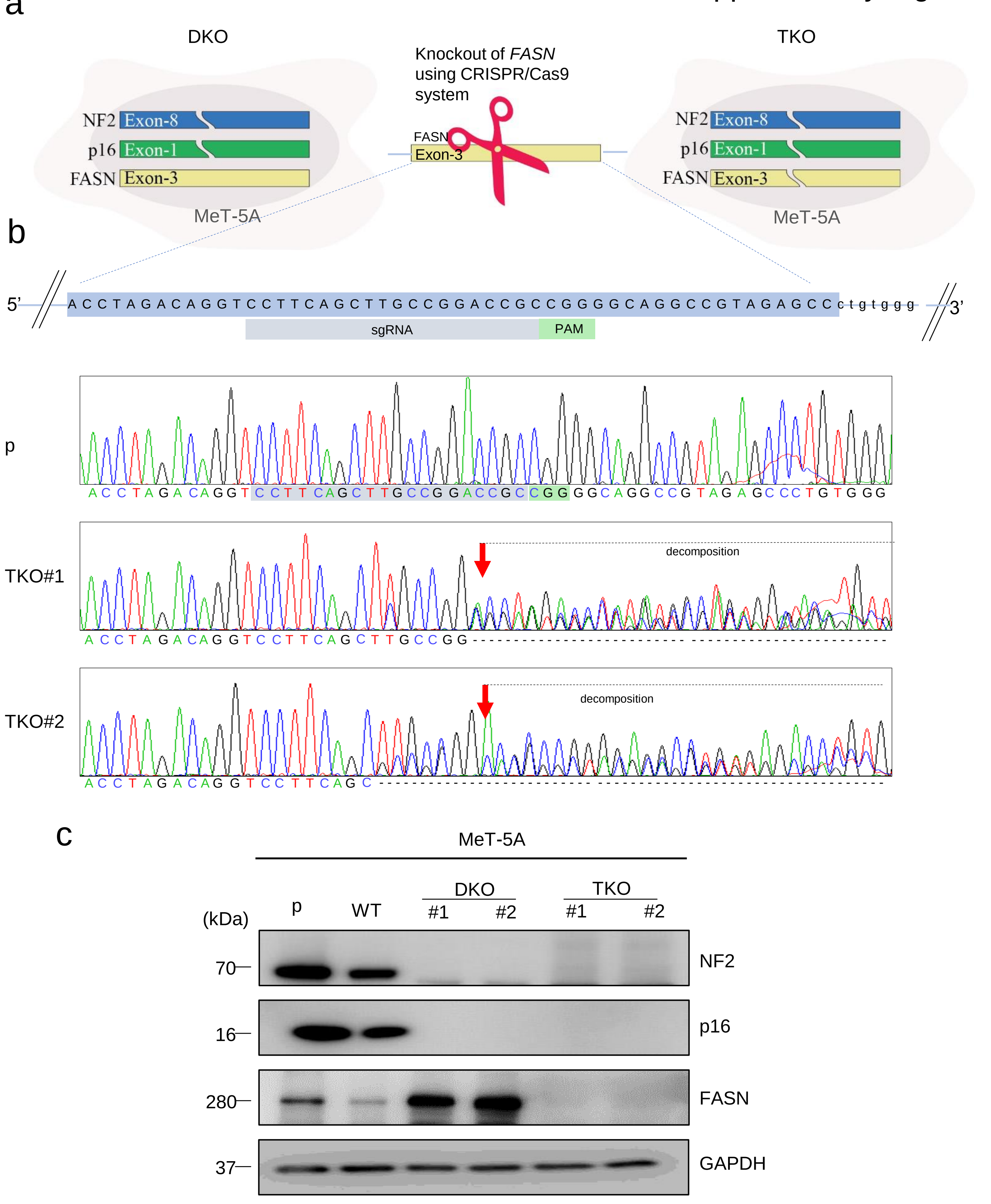


**Supplementary Fig. S2. Generation of *FASN*-knockout MeT-5A-DKO cells (TKO) using the CRISPR/Cas9 system** (a) A specific guide RNA sequence was developed targeting exon 3 of the FASN gene. (b) Genomic sequence comparison between TKO#1, TKO#2, and parental cells was performed to analyze alterations in the FASN locus. (c) Western blot analysis was conducted to assess the protein expressions of NF2, p16, and FASN.

### Supplementary Fig. S3

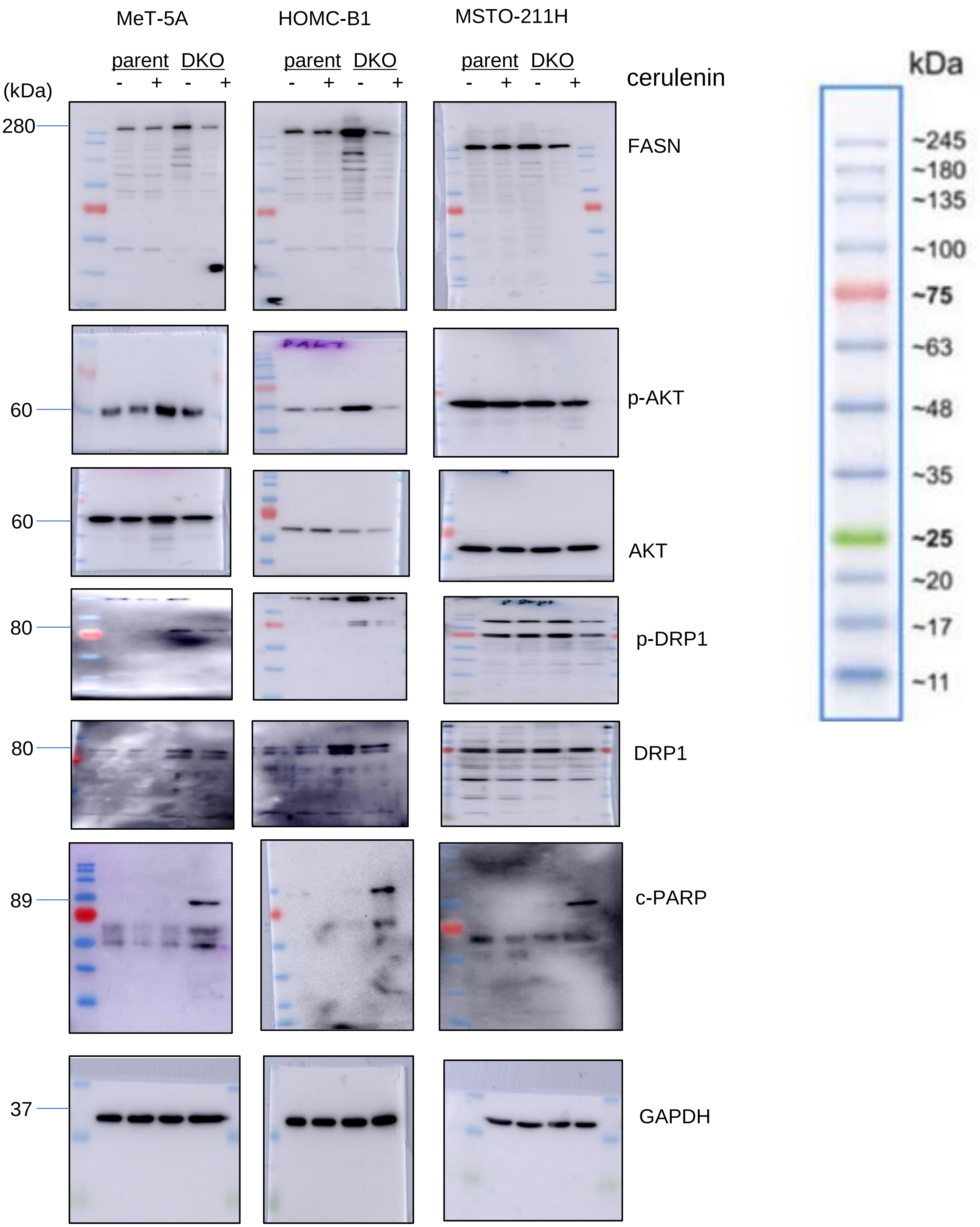


**Supplemental Fig. S3.** Complete and unaltered Western blot image corresponding to Figure 4b

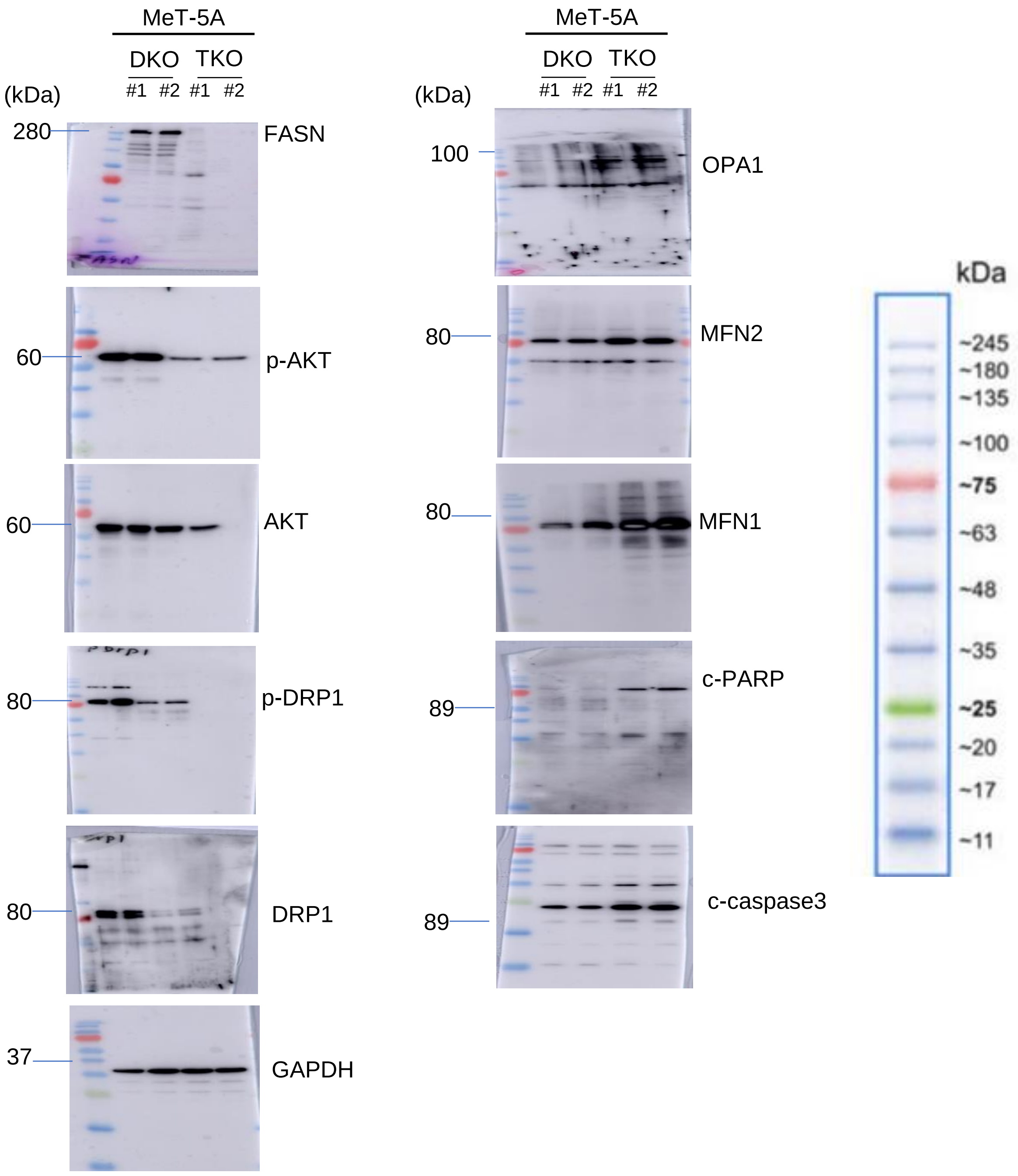


**Supplementary Fig. S4.** Complete and unaltered Western blot image corresponding to Figure 5b

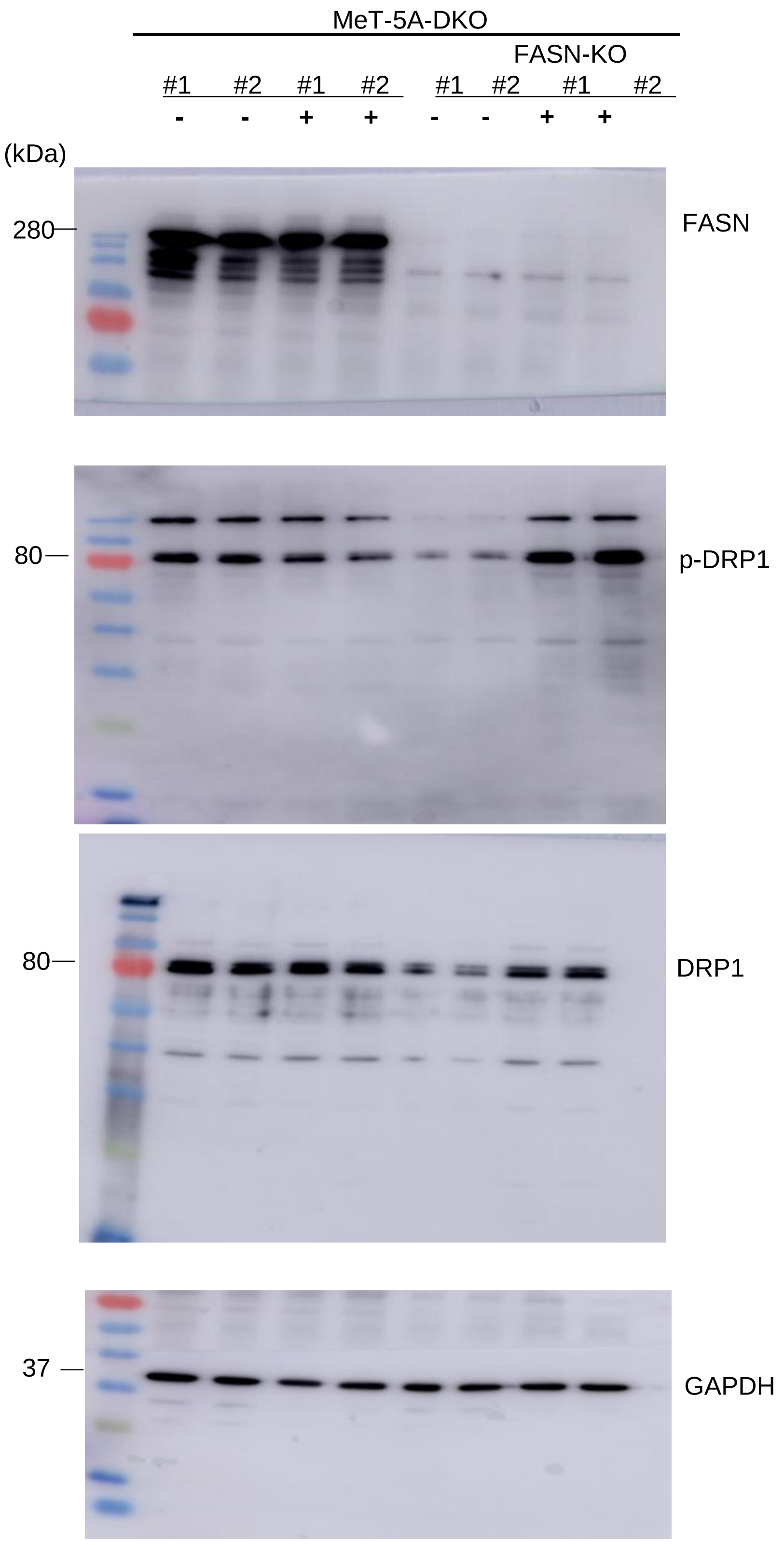


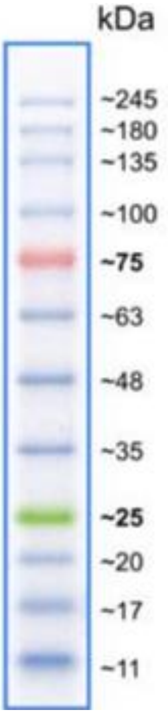


**Supplementary Fig. S5.** Complete and unaltered Western blot image corresponding to Figure 5e

### Supplementary Fig. S6

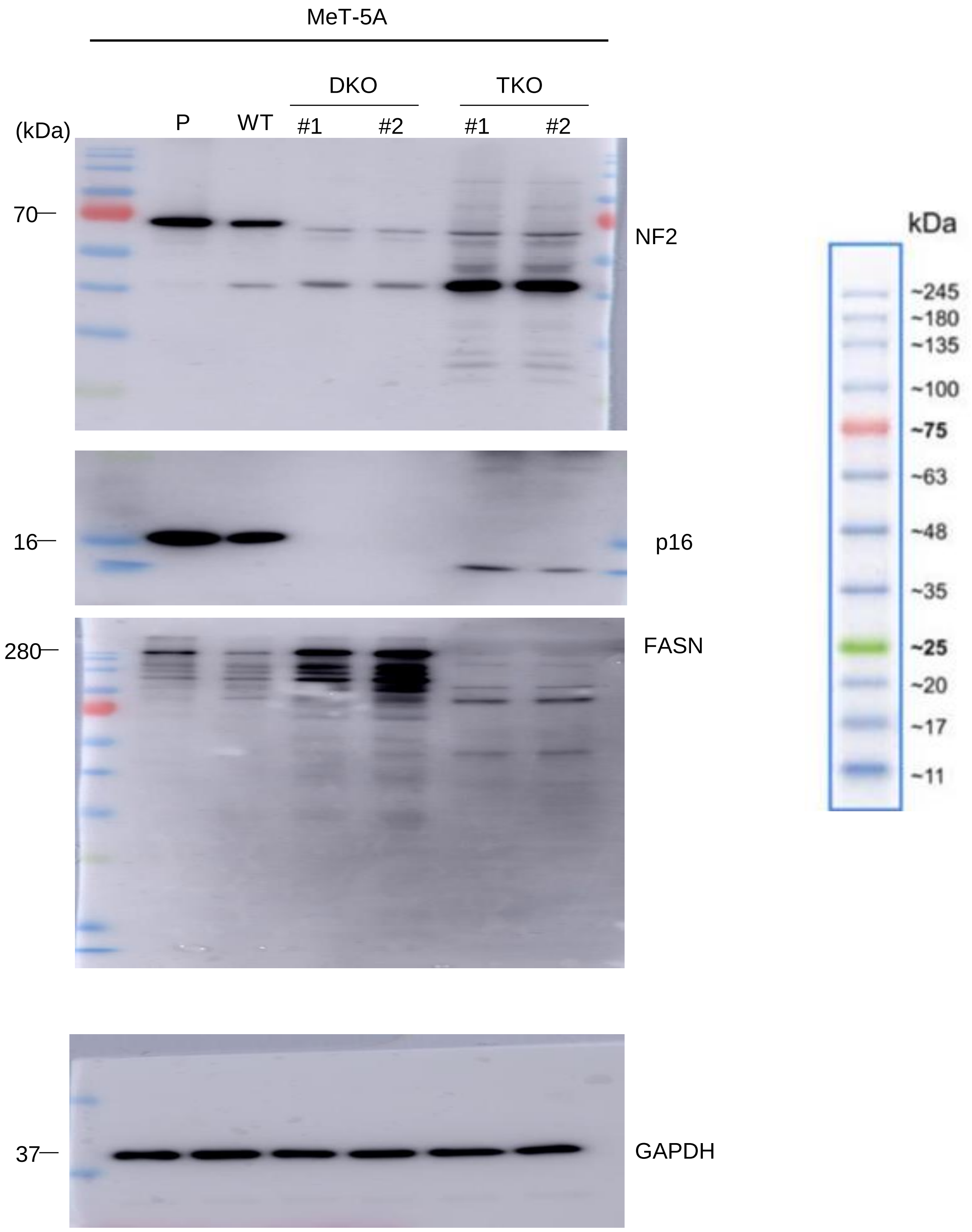


**Supplementary Fig. S6**. Complete and unaltered Western blot image corresponding to Figure S2c

Supplementary Table S1. Effect of 364 compounds on the proliferation of DKO and Parental Met-5A cells.

| No | Category | Compound | MeT-(Parent) cells) | DKO | Relative cell survival |
|---|---|---|---|---|---|
| | | | | (%cells) | DKO-parent (%cells) |
| 1 | blank | DMSO DMSO | 83.88 | 79.41 | -4.47 |
| 2 | antitumor | 5-FU | 67.78 | 67.3 | -0.48 |
| 3 | antitumor | Bestatin | 77.45 | 77.53 | 0.08 |
| 4 | antitumor (DNA) | Bleomycin sulfate | 22.88 | 39.67 | 16.79 |
| 5 | antitumor (DNA) | Cisplatin | 84.6 | 78.42 | -6.18 |
| 6 | antitumor (DHFR) | Methotrexate | 41.39 | 33.72 | -7.67 |
| 7 | antitumor (DNA) | Mitomycin C | 14.35 | 10.95 | -3.4 |
| 8 | antitumor (tubulin) | Vinblastine sulfate | 31.9 | 36.52 | 4.62 |
| 9 | antitumor (tubulin) | Paclitaxel | 68.69 | 47.69 | -21 |
| 10 | antitumor (AR) | Flutamide | 81.14 | 83.62 | 2.48 |
| 11 | antitumor (DNA) | Daunorubicin, HCl | 15.47 | 12.98 | -2.49 |
| 12 | antitumor (DNA) | Doxorubicin, HCl | 22.11 | 12.66 | -9.45 |
| 13 | antitumor (ER) | Tamoxifen, citrate | 37.94 | 61.85 | 23.91 |
| 14 | antitumor (RNA) | Actinomycin D | 20.73 | 15.59 | -5.14 |
| 15 | antitumor (topo I) | Camptothecin | 28.45 | 31.24 | 2.79 |
| 16 | antitumor (topo I/II) | Aclarubicin | 19.01 | 14.48 | -4.53 |
| 17 | antitumor (topo II) | Etoposide (VP-16) | 50.27 | 37.4 | -12.87 |
| 18 | actin filament | Cytochalasin D | 88.95 | 83.63 | -5.32 |
| 19 | adenylcyclase | 2',5'-dideoxyadenosine | 90.95 | 81.74 | -9.21 |
| 20 | AKT | AKT inhibitor | 85.29 | 86.19 | 0.9 |
| 21 | AKT | NL-71-101 | 92.79 | 86.72 | -6.07 |
| 22 | Bcr-Abl | AG957 | 70.76 | 72.44 | 1.68 |
| 23 | caspase | Z-VAD-FMK | 77.32 | 76.53 | -0.79 |
| 24 | CDC2 | Kenpaullone | 55.37 | 59.87 | 4.5 |
| 25 | CDK2 | Purvalanol A | 48.55 | 49.85 | 1.3 |
| 26 | CDK4 | 3-ATA | 60.7 | 85.05 | 24.35 |
| 27 | CDKs | Olomoucine | 81.31 | 80.1 | -1.21 |
| 28 | CKII | TBB | 85.41 | 84.48 | -0.93 |
| 29 | COX-1 | Sulindac sulfide | 85.16 | 80.55 | -4.61 |
| 30 | COX-1 | Valeryl salicylate | 83.25 | 81.66 | -1.59 |

| | | | | | |
|---|---|---|---|---|---|
| 31 | COX-2 | NS-398 | 81.09 | 81.09 | 0 |
| 32 | COX | Sodium salicylate | 87.96 | 82.53 | -5.43 |
| 33 | cyclicphosphodiesterase | Theophylline | 83.93 | 83.6 | -0.33 |
| 34 | DNA methyltransferase | Azacytidine | 82.53 | 84.16 | 1.63 |
| 35 | DNA polymerase | Aphidicolin | 45.28 | 64.34 | 19.06 |
| 36 | EGFR | AG1478 | 84.04 | 82.27 | -1.77 |
| 37 | EGFR, topoII | Genistein | 86.32 | 84.16 | -2.16 |
| 38 | farnesyltransferase | Manumycin A | 32.21 | 56.05 | 23.84 |
| 39 | farnesyltransferase | FTI-276 | 77.9 | 80.88 | 2.98 |
| 40 | Flk-1 | SU1498 | 68.26 | 76.06 | 7.8 |
| 41 | geranylgeranyltransferase I | GGTI-286 | 64.35 | 76.09 | 11.74 |
| 42 | GR | Dexamethasone | 83.75 | 89.43 | 5.68 |
| 43 | GSK-3 | GSK-3 inhibitor II | 77.37 | 88.92 | 11.55 |
| 44 | HDAC | Scriptaid | 23.81 | 26.38 | 2.57 |
| 45 | HDAC | Trichostatin A | 24.19 | 28.82 | 4.63 |
| 46 | HER2 (erbB2/neu), EGFR | AG825 | 82.95 | 76.63 | -6.32 |
| 47 | protein synthesis | Cycloheximide | 34.5 | 32.39 | -2.11 |
| 48 | HMG-CoA reductase | Lovastatin | 39.76 | 66.47 | 26.71 |
| 49 | HSP90 | Radicicol | 25.72 | 27.5 | 1.78 |
| 50 | HSP90 | 17-AAG | 32.49 | 86.54 | 54.05 |
| 51 | IGF-1R | AG1024 | 88.42 | 93.16 | 4.74 |
| 52 | iNOS | 1400W, HCl | 77.91 | 80.65 | 2.74 |
| 53 | iNOS | AMT, HCl | 88.87 | 87.98 | -0.89 |
| 54 | Jak-2 | AG490 | 92.97 | 83.8 | -9.17 |
| 55 | Jak-2 | Cucurbitacin I | 22.65 | 21.21 | -1.44 |
| 56 | JNK | SP600125 | 71.98 | 66.96 | -5.02 |
| 57 | lck (p56), TYK | Damnacanthal | 46.99 | 70.43 | 23.44 |
| 58 | MEK | PD 98059 | 82.41 | 86.45 | 4.04 |
| 59 | MEK | U0126 | 87.63 | 89.84 | 2.21 |
| 60 | methionine aminopeptidase | Fumagillin | 68.08 | 64.61 | -3.47 |
| 61 | MMP | GM 6001 | 84.19 | 77.59 | -6.6 |
| 62 | NF-kB | N-Acetyl-L-cysteine | 84.49 | 78.41 | -6.08 |
| 63 | NOS | Aminoguanidine, HCl | 85.15 | 82.45 | -2.7 |

| | | | | | |
|---|---|---|---|---|---|
| 64 | NOS | L-NMMA | 86.58 | 82.49 | -4.09 |
| 65 | p38 (MAPK) | PD169316 | 80.42 | 72.58 | -7.84 |
| 66 | p38 (MAPK) | SB 203580 | 75.8 | 84.48 | 8.68 |
| 67 | p70 S6K | Rapamycin | 74.45 | 76.92 | 2.47 |
| 68 | PARP | NU1025 | 82.48 | 86.49 | 4.01 |
| 69 | PARP-1 | Benzamide | 83.88 | 81.39 | -2.49 |
| 70 | PC-PLC | D609 | 85.56 | 81.69 | -3.87 |
| 71 | PDE | IBMX | 83.19 | 82.39 | -0.8 |
| 72 | PDE (cAMP) | Ro-20-1724 | 87.14 | 82.91 | -4.23 |
| 73 | PDE (cGMP) | Zaprinast | 84.83 | 90.26 | 5.43 |
| 74 | PDGFR | AG1296 | 63.99 | 50.36 | -13.63 |
| 75 | PI3K | LY294002 | 54.6 | 67.68 | 13.08 |
| 76 | PI3K | Wortmannin | 71.82 | 70.4 | -1.42 |
| 77 | PKA | H-89, HCl | 53.55 | 90.67 | 37.12 |
| 78 | PKC | Bisindolymaleimide I, HCl | 22.65 | 30.62 | 7.97 |
| 79 | PKC, PKA | H-7 | 73.07 | 76.64 | 3.57 |
| 80 | PKC, PKA, PKG, MLCK | Staurosporine | 13.72 | 15.43 | 1.71 |
| 81 | PLA2 | cPLA2inhibitor | 89.03 | 99.16 | 10.13 |
| 82 | PLA2 | OBAA | 61.93 | 61.6 | -0.33 |
| 83 | PP2A | Cantharidin | 84.98 | 89.81 | 4.83 |
| 84 | PP2A | Cytostatin | 81.97 | 90.1 | 8.13 |
| 85 | PP2B/cyclophilin | Cyclosporin A | 94.14 | 85.51 | -8.63 |
| 86 | PP2B/FKBP | FK-506 | 81.08 | 87.59 | 6.51 |
| 87 | proteasome | MG-132 | 25.6 | 27.6 | 2 |
| 88 | proteasome | Lactacystin | 77.62 | 74.84 | -2.78 |
| 89 | ribonucleotide reductase | Hydroxyurea | 88 | 82.66 | -5.34 |
| 90 | ROCK | HA1077 | 76.33 | 78.23 | 1.9 |
| 91 | ROCK | Y27632 | 77.73 | 83.43 | 5.7 |
| 92 | Src, Fyn, Lck | PP1 (analog) | 86.13 | 86.31 | 0.18 |
| 93 | Src, Fyn, Lck | PP-H | 91.84 | 90.59 | -1.25 |
| 94 | tubulin depolymerization | Nocodazole | 37.61 | 46.11 | 8.5 |
| 95 | tyr phosphatase (PTP) | Dephostatin | 89.66 | 74.29 | -15.37 |
| 96 | p53 | Pifithrin-a (cyclic) | 87.73 | 74.52 | -13.21 |

| | | | | | |
|---|---|---|---|---|---|
| 97 | p53 activator | PRIMA-1 | 86.81 | 80.46 | -6.35 |
| 98 | 5α-reductase | Finasteride | 89.4 | 87.09 | -2.31 |
| 99 | aromatase | Aminoglutethimide | 87.32 | 81.82 | -5.5 |
| 100 | aromatase | Formestane | 86.74 | 87.23 | 0.49 |
| 101 | progesterone receptor | Mifepristone | 86.97 | 103.54 | 16.57 |
| 102 | acetyl-CoA carboxylase | TOFA | 86.73 | 87.33 | 0.6 |
| 103 | aminopeptidase A | Amastatin | 86.86 | 75.21 | -11.65 |
| 104 | aminopeptidase M | Actinonin | 85.36 | 77.54 | -7.82 |
| 105 | F1-ATPase | Oligomycin | 24.2 | 48.9 | 24.7 |
| 106 | V-ATPase | Bafilomycin A1 | 13.72 | 21.06 | 7.34 |
| 107 | Bcl-2 | HA 14-1 | 87.19 | 80.28 | -6.91 |
| 108 | Bcl-XL | BH3I-1 | 89.22 | 82.07 | -7.15 |
| 109 | Burton's tyrosine kinase(BTK) | LFM-A13 | 89.58 | 88.99 | -0.59 |
| 110 | Burton's tyrosine kinase(BTK) | Terreic acid | 81.26 | 79.63 | -1.63 |
| 111 | calpain | E-64d | 88.31 | 81.5 | -6.81 |
| 112 | calpain, cathepsin B, L | ALLN | 96.99 | 91.77 | -5.22 |
| 113 | cathepsin B | CA-074 | 90.43 | 82.64 | -7.79 |
| 114 | cathepsin D | Pepstatin A | 92.91 | 80.65 | -12.26 |
| 115 | cathepsin G | Z-GLF-CMK | 25.88 | 23.89 | -1.99 |
| 116 | CCR2 | RS 102895 | 87.42 | 77.5 | -9.92 |
| 117 | CCR3 | SB 328437 | 91.23 | 83.76 | -7.47 |
| 118 | CXCR2 | SB 225002 | 51.98 | 57.67 | 5.69 |
| 119 | CXCR4 | AMD3100 octahydrochloride | 84.59 | 76.12 | -8.47 |
| 120 | Cdc25 | NSC95397 | 29.72 | 28.09 | -1.63 |
| 121 | Cdc25A | SC-αασ9 | 98.79 | 87.06 | -11.73 |
| 122 | Na channel | Amiloride | 92.16 | 82.02 | -10.14 |
| 123 | Na channel | Lidocaine | 94.54 | 81.48 | -13.06 |
| 124 | Na ionophore | Monensin | 35.85 | 56.09 | 20.24 |
| 125 | Na/K ATPase | Ouabain | 30.43 | 30.66 | 0.23 |
| 126 | Na/K/Mg ATPase | Sanguinarine | 13.28 | 9.51 | -3.77 |
| 127 | K channel | Glibenclamide | 86.76 | 77.62 | -9.14 |
| 128 | K channel | Dequalinium | 17.37 | 47.31 | 29.94 |
| 129 | K channel opener | Diazoxide | 92.66 | 81.44 | -11.22 |

| | | | | | |
|---|---|---|---|---|---|
| 130 | K ionophore | Valinomycin | 16.29 | 38.97 | 22.68 |
| 131 | K ionophore | Nigericin | 44.98 | 44.21 | -0.77 |
| 132 | Ca channel | Diltiazem | 91.49 | 76.37 | -15.12 |
| 133 | Ca channel | Nifedipine | 90.67 | 75.4 | -15.27 |
| 134 | Ca channel, MDR | Verapamil | 85.26 | 74.44 | -10.82 |
| 135 | MDR | PGP-4008 | 50.66 | 44.43 | -6.23 |
| 136 | BCRP | Fumitremorgin C | 94.16 | 85.51 | -8.65 |
| 137 | Ca ionophore | A23187 | 13.91 | 11.05 | -2.86 |
| 138 | Ca ionophore | Ionomycin | 17.6 | 18.29 | 0.69 |
| 139 | Ca-ATPase | Thapsigargin | 21.58 | 20.79 | -0.79 |
| 140 | Ca-ATPase | t-Butylhydroquinone (BHQ) | 111.61 | 83.81 | -27.8 |
| 141 | Cl channel | N-phenylanthranilic acid | 88.11 | 81.33 | -6.78 |
| 142 | Cl channel | DIDS | 89.05 | 86.65 | -2.4 |
| 143 | Chk 1 | SB 218078 | 33.28 | 37.96 | 4.68 |
| 144 | Chk 1, 2 | Debromohymenialdisine (DBH) | 95.44 | 83.46 | -11.98 |
| 145 | mitochondrial complex I | Rotenone | 43.96 | 44.95 | 0.99 |
| 146 | mitochondrial complex III | Antimycin A1 | 59.97 | 68.66 | 8.69 |
| 147 | CRM1 | Leptomycin B* | 33.58 | 38.28 | 4.7 |
| 148 | DAG kinase | R59022 | 97.44 | 80.66 | -16.78 |
| 149 | DAG kinase | Dioctanoylglycol | 92.33 | 80.58 | -11.75 |
| 150 | DAG lipase | RHC80267 | 90.58 | 90.56 | -0.02 |
| 151 | DAG acyltransferase (DGAT) | Xanthohumol | 16.18 | 12.03 | -4.15 |
| 152 | fatty acid synthase (FAS) | C75 | 88.8 | 85.6 | -3.2 |
| 153 | FAS | Cerulenin | 80.83 | 32.22 | -48.61 |
| 154 | glycosylation | Tunicamycin | 35.88 | 48.07 | 12.19 |
| 155 | glucosidase I, II | Deoxynojirimycin | 92.97 | 79.06 | -13.91 |
| 156 | a-mannosidase | Swainsonine | 93.91 | 81.24 | -12.67 |
| 157 | guanylate cyclase | LY 83583 | 27.7 | 25.38 | -2.32 |
| 158 | guanylate cyclase | ODQ | 92.94 | 79.78 | -13.16 |
| 159 | HAT | Anacardic acid | 96.63 | 89.41 | -7.22 |
| 160 | HIF | Chetomin | 13.64 | 10.23 | -3.41 |
| 161 | HIF-1a hydroxylase | Dimethyloxalylglycine | 95.11 | 85.94 | -9.17 |

| | | | | | |
|---|---|---|---|---|---|
| 162 | kinesin Eg5 | HR22C16 | 31.11 | 33.13 | 2.02 |
| 163 | kinesin Eg5 | Monastrol | 94.39 | 84.4 | -9.99 |
| 164 | lipoxygenase | Nordihydroguaiaretic acid (NDGA) | 93.32 | 84.93 | -8.39 |
| 165 | 12, 15-lipoxygenase | ETYA | 92.12 | 81.85 | -10.27 |
| 166 | 12-lipoxygenase | Baicalein | 63.86 | 69.08 | 5.22 |
| 167 | Mdm2 | Nutlin-3 | 96.99 | 93.16 | -3.83 |
| 168 | Mdm2 | MDM2 inhibitor | 99.56 | 97.13 | -2.43 |
| 169 | monoamine oxidase | Phenelzine | 91.64 | 84.27 | -7.37 |
| 170 | monoamine oxdase B | Deprenyl | 93.48 | 83.01 | -10.47 |
| 171 | mitochondrial permeability transition pore (MPTP) | Decylubiquinone | 89.55 | 77.16 | -12.39 |
| 172 | MPTP | Ro 5-4864 | 86.69 | 79.14 | -7.55 |
| 173 | MPTP opener | Lonidamine | 92.26 | 84.66 | -7.6 |
| 174 | myosin light chain kinase | ML-7 | 79.86 | 83.86 | 4 |
| 175 | O6-methylguanine-DNA methyltransferase (MGMT) | Benzylguanine | 88.14 | 81.92 | -6.22 |
| 176 | ornithine decarboxylase (ODC) | DFMO | 92.53 | 83.78 | -8.75 |
| 177 | PKG | KT 5823 | 84.31 | 85.1 | 0.79 |
| 178 | PKG | Rp-8-CPT-cGMPS | 78.67 | 75.57 | -3.1 |
| 179 | PPAR-a | MK 886 | 92.33 | 90.17 | -2.16 |
| 180 | PPAR-a activator | Clofibrate | 91.72 | 82.5 | -9.22 |
| 181 | PPAR-g | BADGE | 91.65 | 82.09 | -9.56 |
| 182 | PPAR-g activator | Troglitazone | 98.44 | 93.46 | -4.98 |
| 183 | reverse transcriptase | AZT | 87.72 | 86.04 | -1.68 |
| 184 | reverse transcriptase | Nalidixic acid | 90.65 | 84.02 | -6.63 |
| 185 | RNA polymerase | a-Amanitin | 38.68 | 39.9 | 1.22 |
| 186 | telomerase | MST-312 | 46.36 | 48.58 | 2.22 |
| 187 | telomerase | b-Rubromycin | 70.35 | 63.67 | -6.68 |
| 188 | TGF-b receptor | SB 431542 | 91.74 | 79.48 | -12.26 |
| 189 | spermidine/spermine N1-acetyltransferase (SSAT) activator | N1,N12-Diethylspermine (BESpm) | 85 | 75.91 | -9.09 |
| 190 | sphingosine N-acyltransferase | Fumonisin B1 | 83.55 | 73.19 | -10.36 |
| 191 | AK | ABT-702 | 63.38 | 85.31 | 21.93 |

| | | | | | |
|---|---|---|---|---|---|
| 192 | AKT | Akt Inhibitor IV | 10.64 | 9.23 | -1.41 |
| 193 | AKT | Akt Inhibitor VIII, Isozyme-Selective, Akti-1/2 | 41.49 | 75.65 | 34.16 |
| 194 | AKT | Akt Inhibitor XI | 38.16 | 74.58 | 36.42 |
| 195 | AMPK | compound C | 31.62 | 37.54 | 5.92 |
| 196 | ATM | ATM/ATR kinase inhibitor | 12.04 | 34.41 | 22.37 |
| 197 | ATM | ATM kinase inhibitor | 71.5 | 79.37 | 7.87 |
| 198 | Aurora | Aurora kinase/cdk inhibitor | 47.02 | 45.15 | -1.87 |
| 199 | Aurora | Aurora kinase inhibitor II | 105.48 | 94.36 | -11.12 |
| 200 | Aurora | Aurora kinase inhibitor III | 44.22 | 40.93 | -3.29 |
| 201 | Bcr-abl | AG957 | 74.7 | 50.9 | -23.8 |
| 202 | BTK | LFM-A13 | 94.77 | 88.91 | -5.86 |
| 203 | BTK | Terreic acid | 98.24 | 79.3 | -18.94 |
| 204 | CAMKII | KN-93 | 82.43 | 86.29 | 3.86 |
| 205 | CAMKII | KN-62 | 92.86 | 107.01 | 14.15 |
| 206 | CAMKII | Lavendustin C | 104.8 | 85.3 | -19.5 |
| 207 | CDK | Kenpaullone | 70.81 | 78.26 | 7.45 |
| 208 | CDK | purvalanol A | 55.13 | 66.45 | 11.32 |
| 209 | CDK | Olomoucine | 94.42 | 81.13 | -13.29 |
| 210 | CDK | Alsterpaullone, 2-cyanoethyl | 38.45 | 41.42 | 2.97 |
| 211 | CDK | Cdk1/2 inhibitor III | 35.24 | 33.51 | -1.73 |
| 212 | CDK | Cdk2/9 inhibitor | 42.94 | 42.7 | -0.24 |
| 213 | CDK | NU6102 | 70.93 | 71.24 | 0.31 |
| 214 | CDK | Cdk4 inhibitor | 34.8 | 47.13 | 12.33 |
| 215 | CDK | NSC625987 | 97.12 | 81.57 | -15.55 |
| 216 | Chk | SB218078 | 35.98 | 43.48 | 7.5 |
| 217 | Chk | isogranulatimide | 101 | 83.36 | -17.64 |
| 218 | Chk | Chk2 inhibitor | 94.92 | 78.38 | -16.54 |
| 219 | Chk | Chk2 inhibitor II | 89.89 | 89.96 | 0.07 |
| 220 | CK | Ellagic acid | 92.71 | 86.08 | -6.63 |
| 221 | CK | TBB | 93.15 | 88.39 | -4.76 |
| 222 | CK | DMAT | 73.02 | 53.5 | -19.52 |

| | | | | | |
|---|---|---|---|---|---|
| 223 | CK | D4476 | 93.6 | 75.17 | -18.43 |
| 224 | Clk | TG003 | 91.82 | 79.41 | -12.41 |
| 225 | DGK | Diacylglycerol kinase inhibitor II | 101.57 | 88.69 | -12.88 |
| 226 | DNA-PK | IC60211 | 93.88 | 82.97 | -10.91 |
| 227 | eEF2 | TX-1918 | 57.34 | 54 | -3.34 |
| 228 | EGFR | BPIQ-II | 90.65 | 78.17 | -12.48 |
| 229 | EGFR | AG1478 | 96.4 | 84.96 | -11.44 |
| 230 | EGFR | AG490 | 94.42 | 78.05 | -16.37 |
| 231 | FGFR | SU4984 | 90.88 | 75.55 | -15.33 |
| 232 | FGFR | SU5402 | 78.37 | 76.22 | -2.15 |
| 233 | Flt-3 | Flt-3 Inhibitor | 56.82 | 46.39 | -10.43 |
| 234 | Fms | cFMS Receptor Tyrosine Kinase Inhibitor | 95.1 | 80.51 | -14.59 |
| 235 | Fyn | SU6656 | 66.99 | 58.24 | -8.75 |
| 236 | GSK | GSK-3 inhibitor IX | 93.93 | 85.08 | -8.85 |
| 237 | GSK | 1-Azakenpaullone | 59.77 | 56.2 | -3.57 |
| 238 | GSK | indirubin-3'-monoxime | 59.77 | 59.79 | 0.02 |
| 239 | HER2 | AG825 | 102.58 | 80.18 | -22.4 |
| 240 | IGF-IR | AG1024 | 105.15 | 90.71 | -14.44 |
| 241 | IGF-IR | AGL 2263 | 106.7 | 94.08 | -12.62 |
| 242 | IKK | BMS-345541 | 37.63 | 45.61 | 7.98 |
| 243 | IKK | IKK-2 inhibitor VI | 42.36 | 41.09 | -1.27 |
| 244 | IRAK | IRAK-1/4 inhibitor | 95.26 | 83.23 | -12.03 |
| 245 | Jak | JAK Inhibitor I | 105.18 | 90.11 | -15.07 |
| 246 | Jak | JAK3 Inhibitor VI | 104.36 | 87.24 | -17.12 |
| 247 | JNK | SP600125 | 91.26 | 73.77 | -17.49 |
| 248 | JNK | JNK inhibitor VIII | 95.02 | 87.81 | -7.21 |
| 249 | Lck | Damnacanthal | 83.65 | 71.4 | -12.25 |
| 250 | Lck | PP2 | 66.73 | 65.77 | -0.96 |
| 251 | MAPK | ERK inhibitor II | 85.95 | 71.61 | -14.34 |
| 252 | MEK | PD98059 | 96.2 | 81.87 | -14.33 |
| 253 | MEK | U-0126 | 99.02 | 85.82 | -13.2 |
| 254 | MEK | MEK inhibitor I | 93.22 | 83.18 | -10.04 |

| | | | | | |
|---|---|---|---|---|---|
| 255 | Met | SU11274 | 108.06 | 93.19 | -14.87 |
| 256 | MLCK | ML-7 | 83.73 | 90.68 | 6.95 |
| 257 | p38 | SB202190 | 86.58 | 79.73 | -6.85 |
| 258 | p38 | SB239063 | 89.51 | 73.47 | -16.04 |
| 259 | PDGFR | AG1296 | 88.79 | 60.96 | -27.83 |
| 260 | PDGFR | SU11652 | 11.55 | 11.92 | 0.37 |
| 261 | PDGFR | PDGF receptor tyrosine kinase inhibitor V | 103.62 | 85.62 | -18 |
| 262 | PDGFR | PDGF receptor tyrosine kinase inhibitor IV | 51.9 | 46.3 | -5.6 |
| 263 | PI3K | LY-294002 | 105.91 | 90.74 | -15.17 |
| 264 | PI3K | Wortmannin | 103.19 | 91.01 | -12.18 |
| 265 | PKA | H-89 | 104.39 | 88.54 | -15.85 |
| 266 | PKA | 4-cyano-3-methylisoquinoline | 104.92 | 92.47 | -12.45 |
| 267 | PKC | Bisindolymaleimide I, HCl | 32.52 | 40.55 | 8.03 |
| 268 | PKC | Go7874 | 11.4 | 9.11 | -2.29 |
| 269 | PKG | Rp-8-CPT-cGMPS | 101.8 | 91.2 | -10.6 |
| 270 | PKG | KT5823 | 78.59 | 58.66 | -19.93 |
| 271 | PKR | PKR inhibitor | 34.55 | 43.32 | 8.77 |
| 272 | Raf | RAF1 kinase inhibitor I | 97.5 | 87.9 | -9.6 |
| 273 | Raf | ZM 336372 | 95.74 | 83.42 | -12.32 |
| 274 | ROCK | H-1152 | 68.57 | 70.61 | 2.04 |
| 275 | ROCK | Y-27632 | 90.57 | 82.91 | -7.66 |
| 276 | Hsp90 | radicicol | 30.18 | 33.07 | 2.89 |
| 277 | Src | PP1 analog | 101.33 | 91.58 | -9.75 |
| 278 | Syk | Syk inhibitor | 62.99 | 70.18 | 7.19 |
| 279 | TGF-βRI | SB431542 | 96.65 | 87.58 | -9.07 |
| 280 | TGF-βRI | TGF-b RI kinase inhibitor II | 93.76 | 79.2 | -14.56 |
| 281 | Tpl2 | Tpl2 kinase inhibitor | 93.4 | 83.49 | -9.91 |
| 282 | TrKA | TrkA inhibitor | 86.07 | 67.63 | -18.44 |
| 283 | VEGFR | VEGFR receptor tyrosine kinase inhibitor II | 74.39 | 78.41 | 4.02 |
| 284 | VEGFR | VEGF recptor 2 kinase inhibitor I | 90.78 | 81.43 | -9.35 |
| 285 | VEGFR | SU1498 | 93.22 | 85.17 | -8.05 |

| | | | | | |
|---|---|---|---|---|---|
| 286 | Bcr-Abl | nilotinib | 96.86 | 89.27 | -7.59 |
| 287 | Multi-kinases | sorafenib | 15.45 | 32.64 | 17.19 |
| 288 | mTOR | temsirolimus | 88.41 | 75.8 | -12.61 |
| 289 | EGFR/Her2 | lapatinib | 68.01 | 74.57 | 6.56 |
| 290 | Bcr-Abl/Kit | imatinib mesylate | 95.2 | 95.14 | -0.06 |
| 291 | Multi-kinases | sunitinib malate | 24.38 | 33.3 | 8.92 |
| 292 | EGFR | gefitinib | 94.49 | 98.76 | 4.27 |
| 293 | HDAC | vorinostat | 35.69 | 32.84 | -2.85 |
| 294 | EGFR | erlotinib | 93.7 | 80.7 | -13 |
| 295 | Proteasome | bortezomib | 24.04 | 26.54 | 2.5 |
| 296 | Bcr-Abl/Src | dasatinib | 42.38 | 65.16 | 22.78 |
| 297 | mTOR | everolimus | 72.72 | 73.62 | 0.9 |
| 298 | Multi-kinases | pazopanib | 32.89 | 59.92 | 27.03 |
| 299 | Rho/SRF | CCG-1423 | 90.14 | 87.65 | -2.49 |
| 300 | PIM | PIM1/2 Kinase Inhibitor V | 95.34 | 91.16 | -4.18 |
| 301 | PIM | PIM1 Inhibitor II | 110.86 | 107.14 | -3.72 |
| 302 | Hedgehog | AY 9944 | 104.36 | 100.28 | -4.08 |
| 303 | Hedgehog | cyclopamine | 94.36 | 89.93 | -4.43 |
| 304 | Hedgehog | Jervine | 97.14 | 98.08 | 0.94 |
| 305 | STAT3 | WP1066 | 33.25 | 34.77 | 1.52 |
| 306 | STAT3 | 5,15-DPP | 88.79 | 83.88 | -4.91 |
| 307 | Wnt | IWP-2 | 102.69 | 100.72 | -1.97 |
| 308 | Wnt | IWR-1-endo | 93.27 | 97.23 | 3.96 |
| 309 | Wnt | FH535 | 92.25 | 98.97 | 6.72 |
| 310 | Notch | DAPT | 98.69 | 88.69 | -10 |
| 311 | tankyrase-selective PARP | XAV939 | 90.63 | 86.91 | -3.72 |
| 312 | pan-PARP | PJ-34 | 88.49 | 79.14 | -9.35 |
| 313 | PARP-1/2-selective | Olaparib | 61.55 | 66.34 | 4.79 |

| | | | | | |
|---|---|---|---|---|---|
| 314 | antipsychotic drug | chlorpromazine hydrochloride | 45.16 | 95.67 | 50.51 |
| 315 | depression treatment | desipramine hydrochloride | 92.68 | 94.31 | 1.63 |
| 316 | golgi inhibitor | brefeldin A | 33.68 | 36.48 | 2.8 |
| 317 | stress inducer | anisomycin | 35.27 | 34.64 | -0.63 |
| 318 | thalidomide family | thalidomide | 102.84 | 92.53 | -10.31 |
| 319 | thalidomide family | lenalidomide | 103.66 | 92.96 | -10.7 |
| 320 | retinoids | tretinoin | 96.56 | 93.77 | -2.79 |
| 321 | retinoids | tamibarotene | 96.98 | 104.8 | 7.82 |
| 322 | DNA alkylation | temozolomide | 100.07 | 91.16 | -8.91 |
| 323 | EML4-ALK | crizotinib | 13.15 | 18.46 | 5.31 |
| 324 | mTOR | Torkinib | 106.02 | 93.07 | -12.95 |
| 325 | lipase | orlistat | 58.79 | 84.12 | 25.33 |
| 326 | AR | MDV3100 | 101.82 | 97.42 | -4.4 |
| 327 | caspase activator | PAC-1 | 39 | 41.09 | 2.09 |
| 328 | blc-2 | ABT-737 | 45.67 | 42.23 | -3.44 |
| 329 | G9a | UNC0638 | 20.22 | 43.22 | 23 |
| 330 | G9a | BIX01294 | 30.08 | 73.82 | 43.74 |
| 331 | LSD1 | S2101 (LSD1 inhibitor II) | 97.78 | 90.24 | -7.54 |
| 332 | PRMT1 | AMI-1 | 98.38 | 89.47 | -8.91 |
| 333 | p300 | C646 | 102.23 | 93.28 | -8.95 |
| 334 | SIRT1 | SIRT1 inhibitor III | 79.64 | 68 | -11.64 |
| 335 | SIRT1/2 | Tenovin-6 | 27.2 | 58.34 | 31.14 |
| 336 | HDAC8 | PCI-34051 | 79.4 | 69.31 | -10.09 |
| 337 | BRD4 bromodomain | (+)-JQ1 | 41.9 | 54.14 | 12.24 |
| 338 | Telomerase | TMPyP4 | 96.09 | 78.94 | -17.15 |
| 339 | PARP | BSI-201 (Iniparib) | 110.67 | 95.8 | -14.87 |
| 340 | PARP | ABT-888 (Veliparib) | 93.94 | 83.94 | -10 |
| 341 | PARP | AG014699 (Rucaparib) | 73.51 | 72.26 | -1.25 |
| 342 | PARP | MK-4827 (Niraparib) | 65.98 | 55.14 | -10.84 |

| | | | | | |
|---|---|---|---|---|---|
| 343 | Aurora | ENMD-2076 | 106.38 | 102.53 | -3.85 |
| 344 | Aurora | MLN8237 | 98.21 | 85.92 | -12.29 |
| 345 | Survivin | YM155 | 12.37 | 9 | -3.37 |
| 346 | PDK1 | OSU-03012 | 88.14 | 99.61 | 11.47 |
| 347 | IGF-IR | OSI-906 | 68.77 | 66.67 | -2.1 |
| 348 | c-Met | PF-04217903 | 91.84 | 90.92 | -0.92 |
| 349 | DNMT | Decitabine | 98.16 | 87.74 | -10.42 |
| 350 | Multi-kinases | Vandetanib | 44.5 | 62.42 | 17.92 |
| 351 | Multi-kinases | Axitinib | 62.33 | 69.44 | 7.11 |
| 352 | BRAF | Vemurafenib | 67.58 | 89.01 | 21.43 |
| 353 | JAK | Ruxolitinib | 65.06 | 87.18 | 22.12 |
| 354 | Hedgehog | Vismodegib | 101.13 | 101.94 | 0.81 |
| 355 | GLI1 | Gant61 | 99.4 | 112.46 | 13.06 |
| 356 | FGFR | PD173074 | 40.32 | 54.49 | 14.17 |
| 357 | ALK | A83-01 | 91.87 | 84.43 | -7.44 |
| 358 | GSK-3 | BIO | 75.72 | 69 | -6.72 |
| 359 | GSK-3 | TWS119 | 34.86 | 40.13 | 5.27 |
| 360 | GSK-3 | CT99021 | 86.27 | 106.9 | 20.63 |
| 361 | TGFb-R | LY2157299 | 95.26 | 88.78 | -6.48 |
| 362 | TGFb-R | SD208 | 93.07 | 96.47 | 3.4 |
| 363 | ALK | LDN193189 | 13.36 | 12.55 | -0.81 |
| 364 | ROCK | Thiazovivin | 87.72 | 80.37 | -7.35 |

Supplementary Table S2. Summary of expression status immunohistochemistry in this study

| Cases No. | Age | Sex | Pathology diagnosis | subtype | Intensity | | |
|---|---|---|---|---|---|---|---|
| | | | | | NF2 | p16 | FASN |
| 1 | 64 | M | Malignant mesothelioma of chest wall | S | 2+ | 2+ | 2+ |
| 2 | 48 | M | Malignant mesothelioma | S | 1+ | 1+ | 0 |
| 3 | 58 | F | Malignant mesothelioma | B | 0 | 0 | 2+ |
| 4 | 49 | F | Malignant mesothelioma | S | 0 | 0 | 1+ |
| 5 | 22 | M | Malignant mesothelioma | E | 0 | 0 | 3+ |
| 6 | 49 | M | Malignant mesothelioma | E | 0 | 0 | 1+ |
| 7 | 32 | M | Malignant mesothelioma | E | 2+ | 2+ | 2+ |
| 8 | 29 | M | Malignant mesothelioma with necrosis | E | 2+ | 0 | 3+ |
| 9 | 31 | F | Malignant mesothelioma | S | 2+ | 3+ | 3+ |
| 10 | 70 | F | Malignant mesothelioma | E | 0 | 0 | 2+ |
| 11 | 47 | M | Malignant mesothelioma | S | 2+ | 1+ | 0 |
| 12 | 60 | M | Malignant mesothelioma | E | 1+ | 3+ | 0 |
| 13 | 46 | M | Malignant mesothelioma | E | 1+ | 2+ | 0 |
| 14 | 35 | M | Malignant mesothelioma | E | 1+ | 1+ | 0 |
| 15 | 83 | M | Malignant mesothelioma | B | 2+ | 1+ | 0 |
| 16 | 56 | M | Malignant mesothelioma of mediastinal pleura | S | 1+ | 2+ | 0 |
| 17 | 67 | F | Malignant mesothelioma | E | 3+ | 2+ | 1+ |
| 18 | 40 | F | Malignant mesothelioma of left cardiac atrium | S | 1+ | 1+ | 0 |
| 19 | 50 | F | Malignant mesothelioma of left cardiac atrium | E | 2+ | 3+ | 0 |
| 20 | 43 | F | Malignant mesothelioma of pericardium | E | 3+ | 1+ | 2+ |
| 21 | 43 | F | Malignant mesothelioma of pericardium | E | 3+ | 0 | 2+ |
| 22 | 50 | F | Malignant mesothelioma of pericardium | E | 2+ | 0 | 2+ |

| | | | | | | | |
|---|---|---|---|---|---|---|---|
| 23 | 57 | M | Malignant mesothelioma | E | 0 | 0 | 2+ |
| 24 | 29 | M | Malignant mesothelioma | E | 2+ | 1+ | 0 |
| 25 | 71 | M | Malignant mesothelioma | E | 0 | 0 | 2+ |
| 26 | 63 | F | Malignant mesothelioma | E | 2+ | 0 | 2+ |
| 27 | 48 | F | Malignant mesothelioma | S | 1+ | 2+ | 0 |
| 28 | 60 | F | Malignant mesothelioma | E | 2+ | 2+ | 0 |
| 29 | 60 | F | Malignant mesothelioma | E | 2+ | 1+ | 2+ |
| 30 | 33 | F | Malignant mesothelioma | E | 2+ | 1+ | 0 |
| 31 | 47 | M | Malignant mesothelioma of mesocolon | E | 0 | 0 | 2+ |
| 32 | 44 | F | Malignant mesothelioma of mesocolon | S | 1+ | 1+ | 0 |
| 33 | 60 | M | Malignant mesothelioma | E | 1 | 3+ | 0 |
| 34 | 60 | M | Malignant mesothelioma | E | 0 | 0 | 1+ |
| 35 | 76 | F | Malignant mesothelioma of hypogastrium | E | 2+ | 3+ | 2+ |
| 36 | 78 | M | Malignant mesothelioma with necrosis of hypogastrium | E | 2+ | 3+ | 0 |
| 37 | 41 | F | Malignant mesothelioma of hypogastrium | S | 1+ | 2+ | 0 |
| 38 | 28 | M | Malignant mesothelioma of retroperitoneum | B | 0 | 0 | 1+ |
| 39 | 5 | F | Malignant mesothelioma of epigastrium | E | 0 | 0 | 1+ |
| 40 | 56 | F | Malignant mesothelioma of mesocolon | E | 2+ | 0 | 1+ |
| 41 | 45 | M | Malignant mesothelioma | E | 0 | 0 | 1+ |
| 42 | 69 | M | Malignant mesothelioma | E | 2+ | 3+ | 2+ |
| 43 | 78 | M | Malignant mesothelioma of hypogastrium | E | 0 | 0 | 1+ |
| 44 | 33 | M | Malignant mesothelioma of retroperitoneum | E | 0 | 0 | 1+ |
| 45 | 5 | F | Malignant mesothelioma | E | 0 | 0 | 1+ |

| | | | | | | | |
|---|---|---|---|---|---|---|---|
| 46 | 15 | F | Mesothelium | - | 2+ | 2+ | 0 |
| 47 | 19 | F | Mesothelium | - | 2+ | 1+ | 0 |

Epithelioid (E), Sarcomatous (S), Biphasic (B) FASN expression: Epithelioid 23 out of 32, Sarcomatous 3 out of 10, Biphasic 2 out of 3, Each intensity of the positive signal for NF2, 16 and FASN were evaluated by two investigators, scoring staining intensity as strong (3+), moderation (2+), weak (1+), or negative (0).

Supplementary Table S3: Blood chemistry in vehicle control mice and Cerulenin-treated mice)

| Test | Unit | Control (Mean ± SD) | Cerulenin- treated (20mg/kg, 6 times injection) (Mean ± SD) |
|---|---|---|---|
| TP | g/dL | 4.20 ± 0.04 | 4.02 ± 0.05 |
| ALB | g/dL | 2.84 ± 0.02 | 2.78 ± 0.04 |
| BUN | mg/dL | 21.62 ± 0.54 | 21.74 ± 1.28 |
| CRE | mg/dL | 0.12 ± 0.01 | 0.12 ± 0.01 |
| Na | mEq/L | 149.80 ± 0.80 | 152.00 ± 0.75 |
| K | mEq/L | 5.30 ± 0.18 | 4.40 ± 0.13 |
| Cl | mEq/L | 110.40 ± 0.40 | 114.20 ± 0.33 |
| Ca | mg/dL | 8.28 ± 0.31 | 8.02 ± 0.29 |
| IP | mg/dL | 12.52 ± 0.97 | 13.72 ± 0.52 |
| AST | IU/L | 48.60 ± 2.66 | 42.20 ± 1.78 |
| ALT | IU/L | 26.60 ± 2.25 | 23.60 ± 1.12 |
| LDH | IU/L | 148.60 ± 13.04 | 118.40 ± 5.25 |
| AMY | IU/L | 2511.20 ± 122.93 | 2034.60 ± 71.79 |
| γ-GT | IU/L | 3＞ | 3＞ |
| T-CHO | mg/dL | 76.40 ± 2.89 | 73.60± 2.92 |
| TG | mg/dL | 93.40 ± 11.98 | 88.40 ± 3.85 |
| HDL-C | mg/dL | 39.00 ± 1.00 | 39.20 ± 1.71 |
| T-BIL | mg/dL | 0.06 ± 0.004 | 0.04 ± 0.002 |
| GLU | mg/dL | 128.60 ± 18.74 | 177.40 ± 19.00 |

**Abbreviations:** TP, total protein; ALB, albumin; BUN, blood urea nitrogen; CRE, creatinine; Na, sodium; K, potassium; Cl, chloride; Ca, calcium; IP, inorganic phosphorus; AST, aspartate aminotransferase; ALT, alanine aminotransferase; LDH, lactate dehydrogenase; AMY, amylase; γ-GT, gamma-glutamyl transferase; T-CHO, total cholesterol; TG, triglycerides; HDL-C, high-density lipoprotein cholesterol; T-BIL, total bilirubin; GLU, glucose (blood sugar); bw, body weight

Table S4. Antibodies used in this study.

| Molecule | Cat number | Company | Species | Dilution |
|---|---|---|---|---|
| FASN | sc-48357 | santa Cruz | Ms | x 2000 |
| Phospho-AKT | #4060 | CST | Rb | x 1000 |
| Total-AKT | # 4685 | CST | Rb | x 1000 |
| pDRP1 | 12957-1AP | protentech | Rb | x 1000 |
| DRP1 | AP12957-1 | Protentech | Rb | x 1000 |
| NF2 | #4458 | CST | Rb | x 2000 |
| p16 | #18048 | CST | Rb | x 3000 |
| OPA1 | #3136 | CST | Ms | x 3000 |
| Caspase9 | A2636 | AB clonal | Rb | x 3000 |
| caspase3 | #9661 | CST | Rb | x 2000 |
| GAPDH | #2118 | CST | Rb | x 2000 |
| Anti-rabbit IgG-HRP | #7074 | CST | Goat | x 4000 |
| Hoechst 33342 | SW150 | Dojindo | | x 1000 |
| Alexa Fluor Plus 488 | A32723 | Invitrogen | | x 1000 |
| Alexa Fluor® 568 | A11011 | Invitrogen | | x 1000 |
| Mitotracker® | M7512 | Invitrogen | | x 10000 |

CST, Cell Signaling Technology; Rb, rabbit; Ms, mouse
